# The Lower γ Region Ensures Unidirectional Rotation and Torque Generation in the Latter Half of the 80° Substep of F1-ATPase

**DOI:** 10.64898/2026.05.07.723410

**Authors:** Tomo Uchiyama, Hiroshi Ueno, Meghna Sobti, Emily J. Furlong, Simon H. J. Brown, Alastair G. Stewart, Hiroyuki Noji

**Affiliations:** Department of Applied Chemistry, Graduate School of Engineering, University of Tokyo.; Heart Rhythms Division, The Victor Chang Cardiac Research Institute, NSW, Australia; St Vincent’s Clinical School, Faculty of Medicine, UNSW Sydney, NSW, Australia; Division of Biomedical Science and Biochemistry, Research School of Biology, Australian National University, Canberra, ACT, Australia; School of Science, Molecular Horizons, and the Australian Research Council Centre for Cryo-electron Microscopy of Membrane Proteins, University of Wollongong, Wollongong, NSW, Australia; Research Institute of Planetary Health (RIPH), The University of Tokyo.

**Keywords:** ATP synthase, rotary molecular motor, F_1_-ATPase, single-molecule analysis, cryo-electron microscopy, chemo-mechanical coupling

## Abstract

F_1_-ATPase achieves unidirectional rotation of its γ shaft through coordinated conformational cycling of the α_3_β_3_ ring, yet how the shaft itself enforces directionality remains unclear. Here we analyzed an axle-less TF_1_ lacking the lower half of the rotor shaft by combining single-molecule rotation assays with cryo-EM structural analysis under catalysis conditions. Under ATP-saturated conditions, wild-type TF_1_ exhibited only three pauses per turn corresponding to the catalytic dwells, whereas axle-less TF_1_ exhibited six pauses per turn, indicating the presence of an additional intermediate during rotation. This previously unreported intermediate dwell was observed at 40° between the binding and catalytic dwells. High-speed recordings revealed frequent backsteps confined to the 40° transition between the intermediate and catalytic dwells. Analysis of the dwell-time distributions indicated an approximately zero free energy bias between these two states, consistent with thermally driven interconversion. Cryo-EM resolved three corresponding intermediates—binding, intermediate, and catalytic dwells—showing a major β conformational change from 0° to 40°, but minimal β rearrangement between 40° and 80°, suggesting that the β power stroke is not operative in this interval. Analysis of the γ rotational scheme indicates that the driving force changes across the 0–80° step: the 0–40° advance proceeds via a power stroke to an intermediate dwell, whereas the subsequent 40° transition to the catalytic dwell lacks a power stroke and is dominated by thermal fluctuations. Taken together, these findings suggest that the lower γ region contributes to unidirectional rotation and torque generation specifically in the latter half of the 0–80° step, while the initial half can proceed even without contributions from the lower γ region.

## Introduction

F_o_F_1_-ATP synthase (F_o_F_1_) is one of the core enzymes that sustain cellular energy systems. This ubiquitous enzyme is found in the plasma membranes of prokaryotic cells, the thylakoid membranes of chloroplasts, and the inner membranes of mitochondria (1, 2). F_o_F_1_ synthesizes ATP from ADP and inorganic phosphate (Pi) using the electrochemical potential difference generated across biological membranes, referred to as the proton motive force (*pmf*). The enzyme is composed of two rotary molecular motors, F_o_ and F_1_, that are mechanically coupled (3). F_o_ is a membrane-embedded motor that generates torque upon proton translocation driven by *pmf*, whereas F_1_ is the membrane-protruding motor that catalyzes ATP synthesis and hydrolysis. Through mechanical coupling via a shared central stalk connecting the two rotors and a peripheral stalk linking the stators, F_o_F_1_ interconverts the *pmf* and the free energy of ATP hydrolysis: under high *pmf*, F_o_ forcibly rotates F_1_ in the reverse direction to induce ATP synthesis; when *pmf* diminishes, F_1_ hydrolyzes ATP and drives the reverse rotation of F_o_, pumping protons across the membrane.

When isolated from F_o_, F_1_ functions solely as an ATPase and is therefore referred to as F_1_-ATPase. The minimal functional composition of F_1_ as a rotary motor is α_3_β_3_γ, in which the central γ subunit rotates against the α_3_β_3_ stator ring (4). Three α and three β subunits alternate to form a heterohexameric stator ring penetrated by γ. The α_3_β_3_ ring possesses two distinct types of nucleotide-binding interfaces: catalytic and non-catalytic interfaces. At the catalytic interface, most residues essential for ATP hydrolysis reside on β, with a key contribution from an arginine residue on α, termed the arginine finger (5, 6); accordingly, β is referred to as the catalytic subunit. At the non-catalytic interface, the nucleotide-binding site is formed primarily by residues on α, which is therefore termed the non-catalytic subunit. During catalysis, the three β subunits undergo coordinated conformational transitions associated with their catalytic cycle, collectively generating the sequential structural changes that drive unidirectional rotation of γ (7–10).

The rotation features of F_1_ have been extensively investigated by single-molecule rotation assay on the F_1_ from the thermophilic bacterium *Bacillus* PS3 (TF_1_). In F_1_, γ rotates counterclockwise when viewed from the F_o_ side, with unitary steps of 120° consistent with the pseudo-three-fold symmetry of the α_3_β_3_ stator. Each full turn is tightly coupled with three ATP hydrolysis events. Torque generation against viscous friction has been estimated to be approximately 40 pN·nm. The high reversibility and efficiency of energy conversion were also experimentally confirmed (11, 12).

The chemomechanical coupling scheme—how rotation and ATP hydrolysis are coupled—has been largely established; each unitary 120° rotation is subdivided into an 80° substep triggered by ATP binding and a subsequent 40° substep associated with ATP cleavage (13, 14). Accordingly, the dwells preceding the 80° and 40° substeps are termed the binding dwell and catalytic dwell, respectively (15). As a consequence, F_1_ exhibits six distinct dwells per rotation, during which the three β subunits sequentially undergo ATP hydrolysis, yielding three ATP molecules per turn.

For an individual β subunit, the reaction scheme can be described as follows. The orientation of the γ subunit, when the β subunit awaits ATP binding is defined as 0°, corresponding to one of the binding-dwell angles. This β subunit hydrolyzes the bound ATP after the γ subunit rotates by approximately 200°, corresponding to a catalytic dwell. ADP release is proposed to occur during the subsequent rotation between 240° and 320° (9, 16), whereas Pi release is likely to occur at 320° (16, 17).

Previous crystallographic studies predominantly captured F_1_ in the catalytic-dwell state, leaving the binding-dwell state structurally unresolved. Cryo-EM single-particle analysis subsequently captured TF_1_ pausing at the binding dwell, thereby providing the missing structural counterpart to the reaction scheme established by single-molecule rotation studies (18).

Thus, the chemomechanical coupling reaction scheme has been well established; however, the molecular mechanisms underlying torque generation and transmission within the F_1_ motor remain incompletely resolved. So far, substantial effort has been devoted to elucidating the molecular mechanisms for that purpose. Structural studies have identified three major β–γ interaction sites—the hydrophobic sleeve, the β-catch loop, and the β-lever domains—arranged along the γ axle from the αβ N-terminal side (19–21). At the hydrophobic sleeve, residues on α and β surround the tip of γ and are thought to function as a bearing, whereas at the β-lever domains, large conformational changes in the C-terminal domain of β are proposed to generate torque for the γ rotation.

Early studies focused on specific point interactions between β and γ through site-directed mutagenesis; however, these studies did not identify residues that were individually crucial for torque generation (22, 23). Subsequent work therefore employed more extensive mutagenesis approaches, most notably a series of large-scale truncation analyses of the γ subunit performed by the Kinoshita group. These studies showed that even mutants lacking most of the γ axle, retaining only minimal contacts with the β-lever domain, still exhibited unidirectional rotation, albeit at markedly reduced rotation rate and torque (24–26). Importantly, later mutagenesis studies that directly targeted the β-lever domain itself further demonstrated that disruption of β-lever–γ interactions does not abolish unidirectional rotation, but instead leads to quantitative reductions in rotation rate and torque (27). Together, these results indicate that torque generation in F_1_ is mediated by the β–γ interaction as a whole, rather than being transmitted through a few specific interactions. However, it has not yet been systematically examined whether these interaction sites contribute uniformly across the entire rotational cycle of F_1_ or whether specific interactions play dominant roles at particular rotational substeps.

Regarding this issue, Furlong et al. analyzed the structure of the axle-less TF_1_ mutant that Kinosita group developed (28). They resolved the cryo-EM structure of TF_1_(γΔN4C25); 4 and 25 residues of the N- and C-terminal helices of γ were deleted. This mutant lacks the lower half of the γ axle and resultantly loses the interactions at the hydrophobic sleeve and β-catch loop (Fig. 1). The mutant was previously reported to exhibit markedly reduced torque compared with wild-type TF_1_. The cryo-EM analysis demonstrated that a canonical binding-dwell architecture can still be formed even in the absence of the γ axle. However, while this finding established the structural integrity of the binding dwell, it remained unclear how truncation of the γ axle affects subsequent rotational substeps or how it contributes to torque generation.

**Fig. 1.**
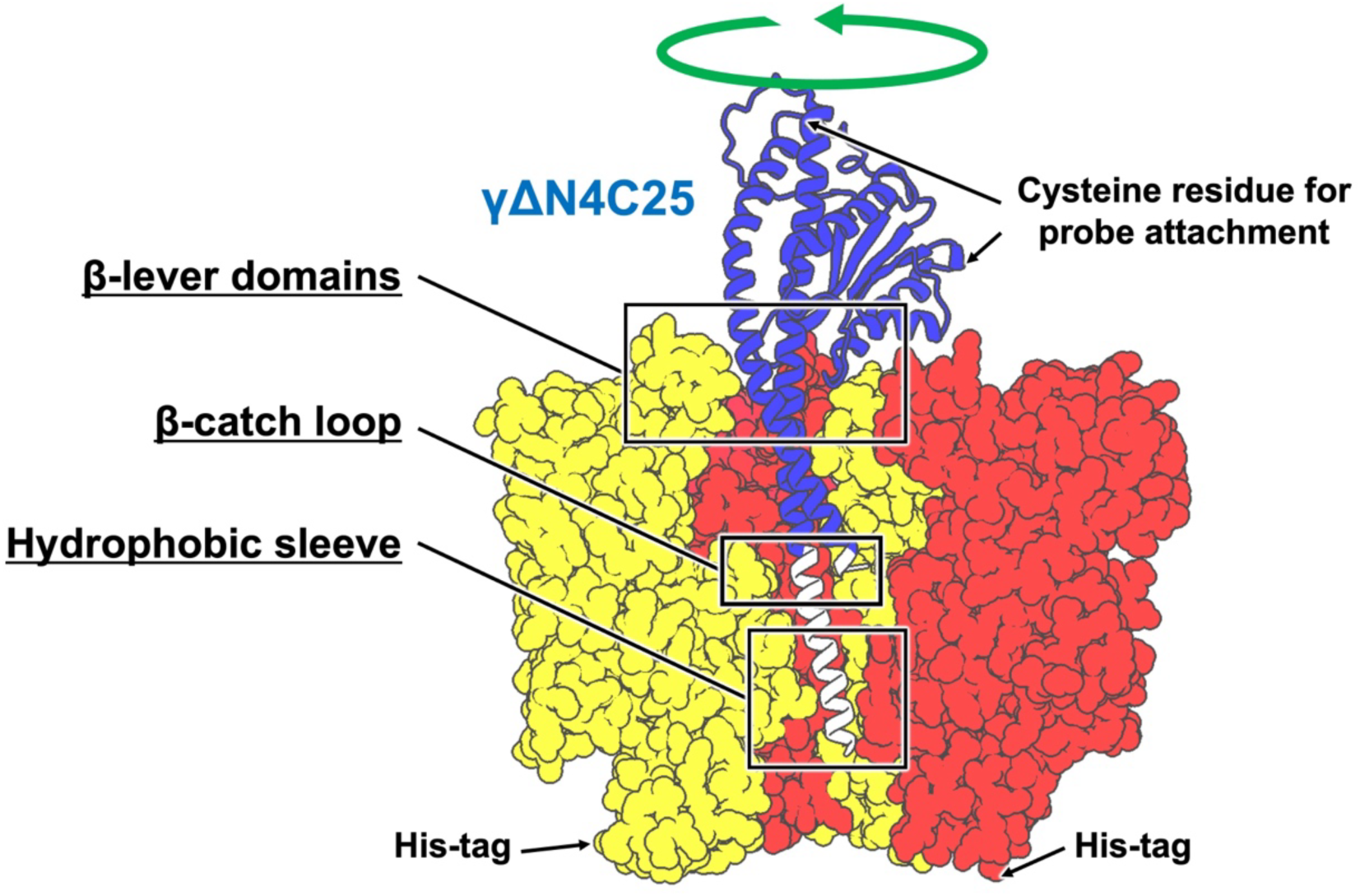
The design of axle-less TF_1_. The structure of TF_1_ with the deleted N-terminal 4 and C-terminal 25 residues of the γ subunit shown in white. The remaining γ region is shown in blue, α in red, and β in yellow. The front αβ pair is removed to show the γ structure. The green arrow indicates the direction of rotation during ATP hydrolysis (counterclockwise as viewed from the membrane side). The black squares indicate the β–γ interaction sites: the hydrophobic sleeve (α280–285, β271–275, γ271–283), the β-catch loop (β312–315, γ262–263), and the β-lever domain (β382–396, γ15–31 and γ243–253) (28).

In this study, we examine how truncation of the γ axle influences the rotational behavior of F_1_, with a particular focus on its effects on individual substeps and associated rotary intermediates during chemomechanical coupling. To this end, we exploited the mutant TF_1_(γΔN4C25) with the deletion of the lower half of the γ axle that hereafter we refer to as axle-less TF_1_ for simplicity. By combining single-molecule rotation assays with cryo-EM structural analysis under catalytic conditions, we determine how loss of specific β–γ interactions alters rotary intermediates and the associated energy landscape.

## Results

### Construct of axle-less TF_1_ for rotation assays

The expression vector encoding axle-less TF_1_ was constructed by introducing the truncation mutation, γ(ΔN4C25) into the wild-type TF_1_ with His-tag at the N-termini of the α and β subunits and two cysteine residues of the γ subunit for the rotation assay according. The expressed protein of axle-less TF_1_ was purified according to the previous report (see Methods for details) (26). Gel filtration chromatography of the purified axle-less TF_1_ protein showed a distinct peak corresponding to the α₃β₃γ complex (Fig. S1A). SDS-PAGE analysis of the peak fraction confirmed the presence of all component subunits, and the γ subunit was shortened as designed (Fig. S1B). The intensity of the γ band was lower than that observed for WT TF_1_, as found in the previous report (28), suggesting that a fraction of the complexes lacked the γ subunit. Nevertheless, because sufficient numbers of rotating molecules were obtained in single-molecule assays, this preparation was judged to be adequate for rotation experiments and was used in the subsequent analyses.

### Rotation kinetics and stepping behavior of axle-less TF_1_

The rotation of axle-less TF_1_ was observed using 40-nm gold colloid as the probe under a laser dark-field microscopy (Fig. 2A). Rotations were recorded at 10,000 frames per second (fps) for WT TF_1_, and at 500–1,000 fps for axle-less TF_1_ depending on [ATP]. The difference in frame rate reflects the limitations in the on-board memory of the high-speed camera; for slowly rotating axle-less TF_1_, the slower frame rates were chosen to optimally balance sufficient temporal resolution with a sufficiently long recording duration to capture a large number of rotational events within a single acquisition. Fig. 2B shows the Michaelis–Menten curves of the rotation rate. The maximum rotation rate (*V*_max_) and Michaelis constant (*K*_m_) were 180 rps and 24 µM for WT TF_1_, and 11 rps and 7.1 µM for axle-less TF_1_, respectively. The *V*_max_ and *K*_m_ values of axle-less TF_1_ were approximately one-tenth and one-third of those for WT TF_1_, respectively. The ATP-binding rate constant, *k*_on_, estimated as 3 × *V*_max_ / *K*_m_, was determined to be 2.3 × 10^7^ M^−1^ s^−1^ for WT TF_1_ and 4.6 × 10^6^ M^−1^ s^−1^ for axle-less TF_1_. The *k*_on_ value of axle-less TF_1_ was approximately one-fifth of that for WT TF_1_. These show that axle-less TF_1_ has evidently slow kinetics with slow ATP binding rate and slow catalytic process—ATP cleavage and/or product release in comparison with the wild-type.

**Fig. 2.**
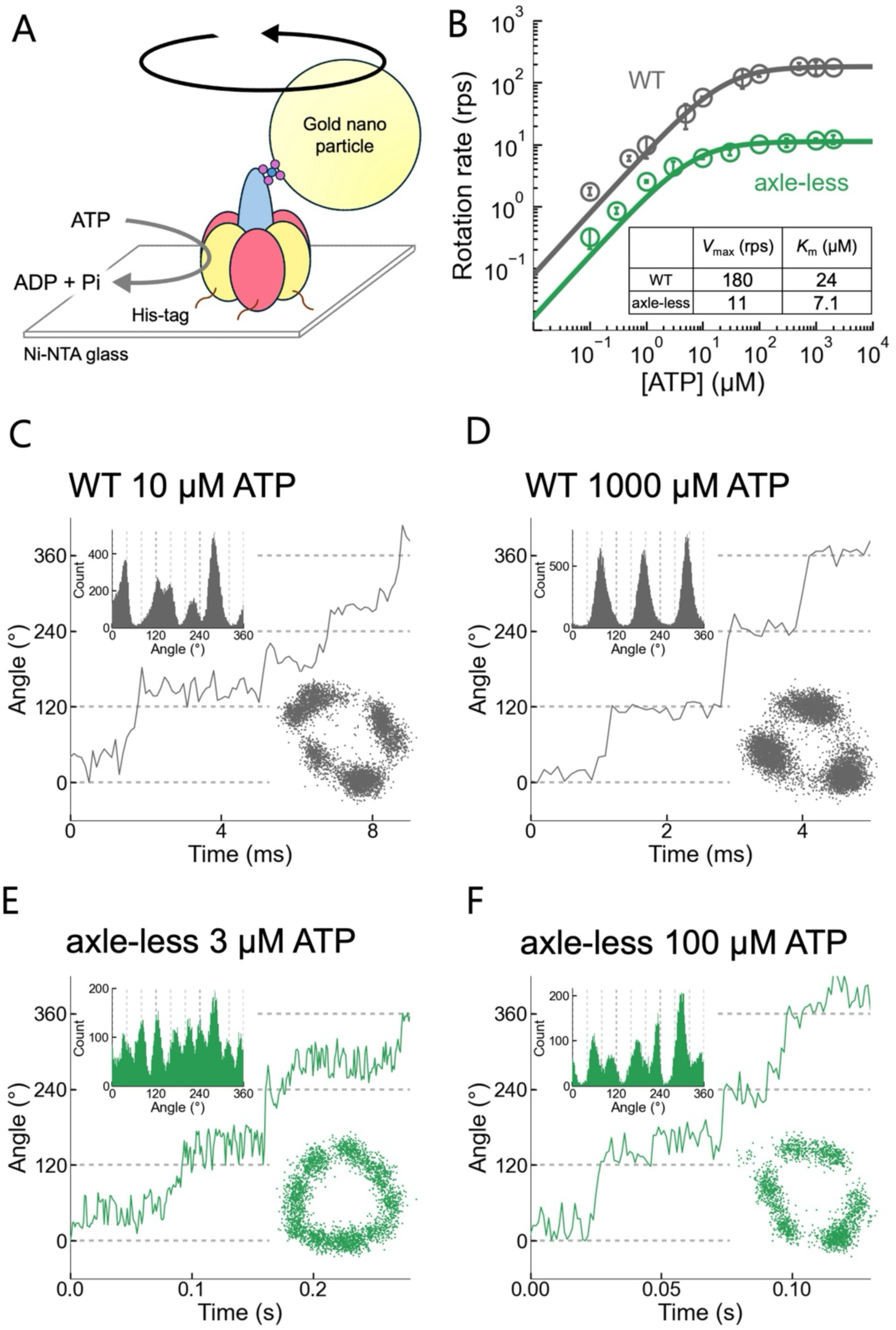
Single-molecule rotation assays of WT TF_1_ and axle-less TF_1_. (A) A schematic image of the single-molecule rotation assay for F_1_. Fixed α_3_β_3_ stator ring was immobilized on a glass surface by His-tag, and the gold colloid probe was attached to the rotor γ subunit through the biotin-streptavidin interaction. (B) [ATP] is plotted against the rotation rate for WT TF_1_ (gray) and axle-less TF_1_ (green). The mean and SD of each data point are shown as circles and error bars, respectively (n = 3–6). Solid lines indicate Michaelis–Menten fits. *V*_max_^WT:^ 180 rps, *K*_m_^WT^: 24 µM. *V*_max_^axle-less^: 11 rps, *K*_m_^axle-less^: 7.1 µM. (C–F) Representative rotation time courses. The x–y plots (right) and angular position histograms (left) are shown as insets. (C, D) Rotation of WT TF_1_ recorded at 10,000 fps. (C) 10 µM ATP. (D) 1000 µM ATP. (E, F) Rotation of axle-less TF_1_ recorded at 1,000 fps. (E) 3 µM ATP. (F) 100 µM ATP.

At low [ATP] far below *K*_m_, axle-less TF_1_ exhibited three distinctive pauses with 120° intervals, which should correspond to the binding dwells. The histogram of dwell time for ATP binding followed a single exponential decay function (Fig. S2). The rate constants determined from the dwell-time histograms yielded *k*_on_ values of 7.7–7.9 × 10^6^, which agrees well with the value estimated from Michaelis–Menten analysis.

Time courses, x–y plots, and angular position histograms demonstrate stepping rotation of WT TF_1_ and axle-less TF_1_ (Fig. 2C–F). At high [ATP] above *K*_m_, WT TF_1_ showed three pauses with 120° intervals (Fig. 2D; Fig. S3B), which correspond to the catalytic dwells. Under the same conditions, axle-less TF_1_ exhibited six characteristic pauses with ∼80° and ∼40° substeps (Fig. 2F; Fig. S3D), indicating the presence of additional pauses not observed in WT TF_1_. Around the *K*_m_ range, WT TF_1_ exhibited six pauses with 80° and 40° substeps (Fig. 2C; Fig. S3A), corresponding to the binding and catalytic dwells. Under the same conditions, axle-less TF_1_ exhibited nine characteristic pauses with ∼40° substeps (Fig. 2E; Fig. S3C). Importantly, three pauses that became prominent only at low [ATP] emerged at angular positions near the midpoint of the ∼80° substeps observed at high [ATP]. Assuming similar angular positions to WT TF_1_, the three pauses following the ∼40° substeps observed at high [ATP] are likely to correspond to the catalytic dwells. Taken together, these results suggest that axle-less TF_1_ exhibits an additional intermediate dwell state between the canonical binding and catalytic dwells that is not observed in WT TF_1_.

Notably, axle-less TF_1_ frequently made backward steps during rotation (Fig. 2E, F; Fig. S3C, D). Although the pauses immediately before and after a backward step were estimated to be in the millisecond range, detailed analysis at 1,000 fps is not practical. We therefore revisit backward steps later using higher-frame-rate recordings.

### Identification of the binding dwell angular positions

To further clarify the angular positions of the binding dwells in axle-less TF_1_, we performed buffer exchange experiments in which [ATP] was switched between a low concentration around the *K*_m_ range and a high concentration above *K*_m_. By comparing the pause angles under the two conditions, we found that six of the nine pauses observed at low [ATP] overlapped with the pauses at high [ATP], whereas three pauses with 120° intervals almost disappeared at high [ATP] (Fig. 3A; Fig. S4). We therefore assigned the three ATP-dependent pauses to the binding dwells. The analysis of the angular difference between a binding dwell position and the next dwell position in the rotational direction revealed a histogram centered at 41 ± 6° (Fig. 3B). Among the six ATP-independent pauses, three are expected to correspond to the catalytic dwells and the other three to an additional dwell state not observed in WT TF_1_.

**Fig. 3.**
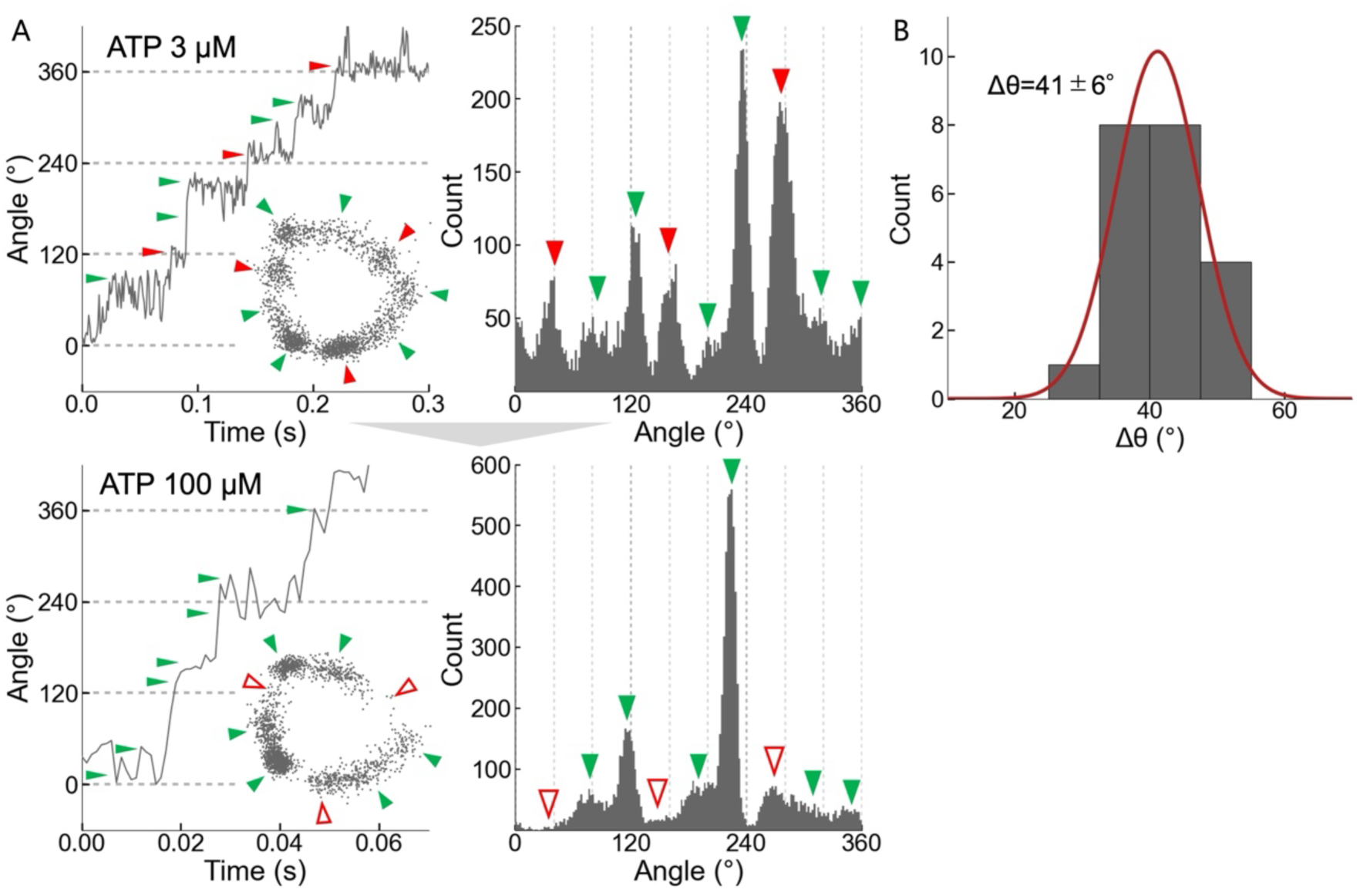
Identification of the binding dwell by buffer exchange. (A) Rotation time courses and angular histograms for the same molecule during a buffer exchange experiment, in which [ATP] was switched between a low concentration around the *K*_m_ range (top) and a high concentration above *K*_m_ (bottom). The x–y plots are shown as insets. Pauses indicated by red arrows were identified as binding dwells. Among the pauses indicated by green arrows, three are inferred to correspond to catalytic dwells and the remaining three to an additional dwell state not observed in WT TF_1_. Videos were recorded at 1,000 fps. (B) Histogram of the angular difference (Δθ), between a binding dwell position and the next dwell position in the direction of rotation. Values are mean ± SD (N = 21, 8 molecules).

### Identification of the catalytic dwell angular positions

ATPγS, a slowly hydrolyzable ATP analog, is known to markedly prolong the catalytic dwell in F_1_-ATPase (13). We used ATPγS to identify the catalytic dwell of axle-less TF_1_. Based on the rotational scheme of WT TF_1_, the catalytic dwell is expected to occur ∼80° ahead of the binding dwell; however, shortening the γ subunit may shift the catalytic dwell positions. The rotation rate of axle-less TF_1_ was measured over a range of [ATPγS] values and fitted with a Michaelis–Menten curve (Fig. S5). The *V*_max_ and *K*_m_ were determined as 2.8 rps and 1.6 µM, respectively. Although *V*_max_ was reduced to approximately one-quarter of that for ATP-driven rotation, this decrease was much smaller than that reported for WT TF_1_, suggesting that the overall turnover of axle-less TF_1_ is not limited solely by ATPγS-sensitive reactions.

We then performed buffer-exchange experiments to switch the substrate from ATP to ATPγS. Both substrates were used at concentrations above *K*_m_, where axle-less TF_1_ exhibited ∼80° and ∼40° substeps (Fig. 4A). In ATPγS buffer, axle-less TF_1_ selectively prolonged the dwell following the ∼40° substep (Fig. 4A; Fig. S6). Based on these observations, we assigned the pauses after the ∼40° substeps as catalytic dwells and the three pauses preceding the ∼40° substeps, i.e., after the ∼80° substeps, as the axle-less TF_1_-specific dwell state, which we termed the “new dwell.” The angular difference between the new dwell and the catalytic dwell was centered at 39° (Fig. 4B). Together with the abovementioned finding that the new dwell is located ∼40° ahead of the binding dwell, these results revealed that axle-less TF_1_ exhibits binding and catalytic dwell states at the same angular positions as WT TF_1_, with an additional dwell located ∼40° between them (Fig. 4C).

**Fig. 4.**
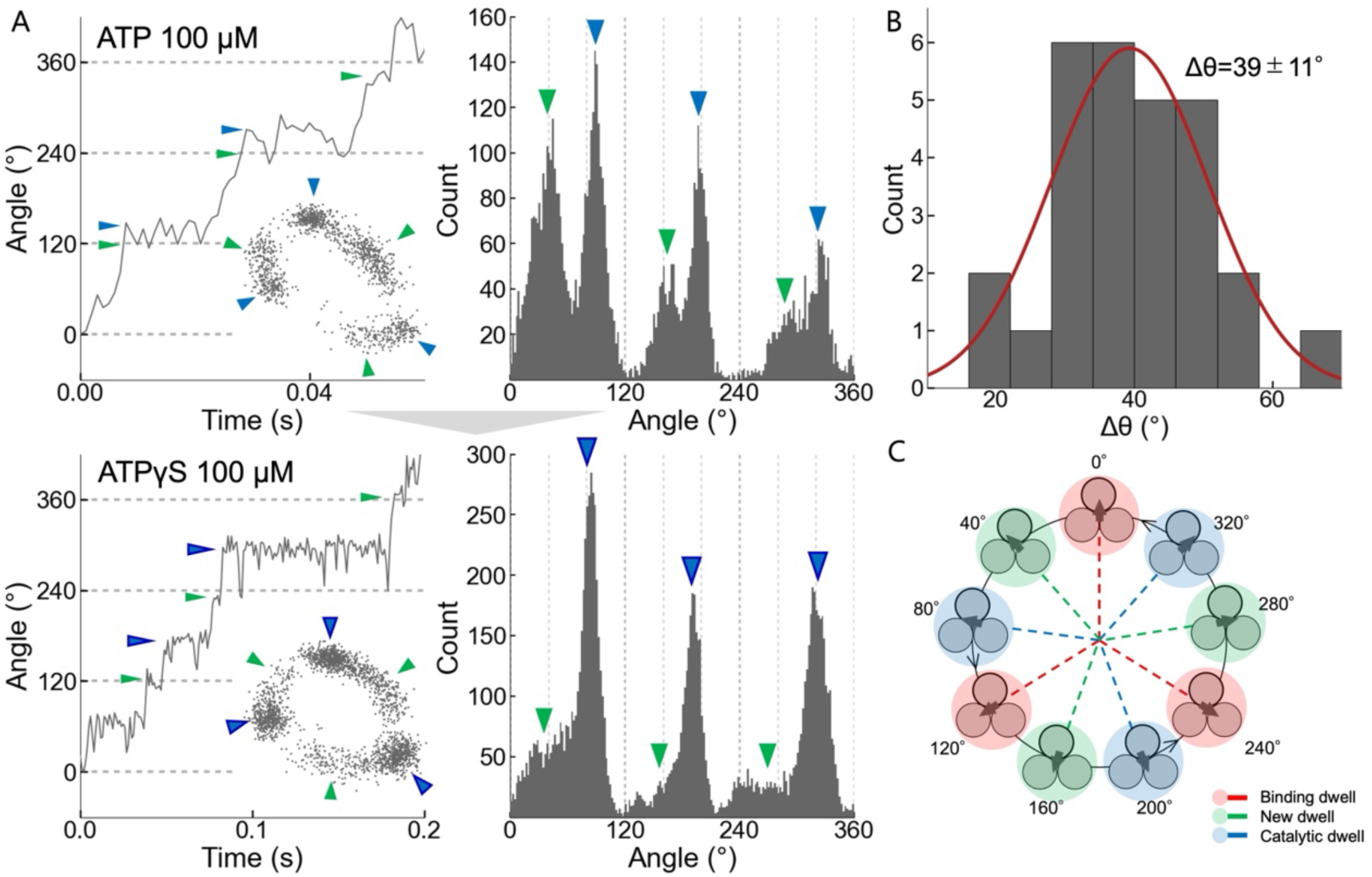
Identification of the catalytic dwell from buffer exchange experiments. (A) Rotation time courses and angular histograms for the same molecule during a buffer exchange experiment, in which the substrate was switched between 100 µM ATP (top) and 100 µM ATPγS (bottom). The x–y plots are shown as insets. Pauses indicated by blue arrows were identified as catalytic dwells. Pauses indicated by green arrows were termed as the “new dwell,” unique to axle-less TF_1_. Videos were recorded at 1,000 fps. (B) Histogram of the angular difference (Δθ), between the catalytic dwell and new dwell positions. Values are mean ± SD (N = 28, 10 molecules). (C) Rotational scheme model of axle-less TF_1_. The three circles represent the three β subunits in the stator ring. The arrow indicates the rotational angular position of the γ subunit. The angular positions of the binding dwell, new dwell, and catalytic dwell are indicated in red, green, and blue, respectively.

### Backward step analysis

For quantitative analysis of backward steps (Fig. 2E, F; Fig. S3C, D), the rotation of axle-less TF_1_ was recorded at 30,000 fps. The ATP concentration was set at 100 µM where axle-less TF_1_ exhibits the new and the catalytic dwells (Fig. 5A). The predominant backward event was a backward step from the catalytic dwell to the new dwell located ∼40° behind. Other backward steps, such as backward steps from the new dwell to the catalytic dwell located ∼80° behind, as well as backward steps larger than 120° from any dwell, were rarely observed. Within our temporal resolution, forward steps from the new dwell always proceeded via the catalytic dwell, and no direct transitions to the next new dwell were observed. Hereafter, we describe the rotational transitions using three types: (i) a forward step from the new dwell to the catalytic dwell located ∼40° ahead, (ii) a forward step from the catalytic dwell to the next new dwell located ∼80° ahead, and (iii) a backward step from the catalytic dwell to the new dwell located ∼40° behind. The reaction scheme and the corresponding rate constants are defined as follows.

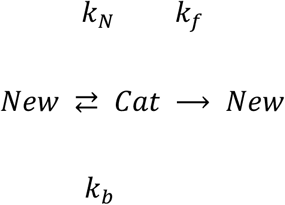

**Fig. 5.**
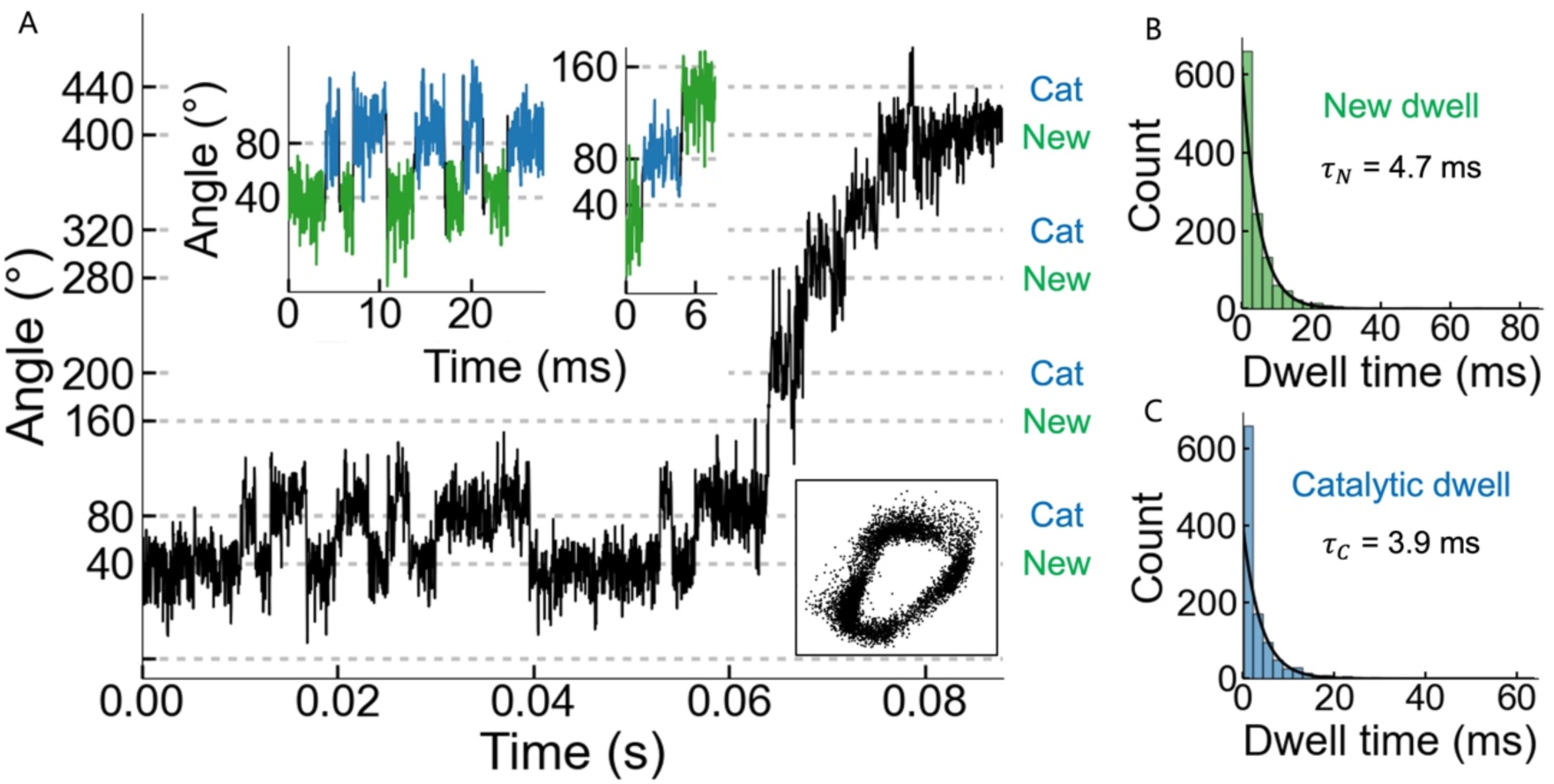
Backward steps between the new dwell and the catalytic dwell at high [ATP]. (A) Representative time course showing a typical backward step during rotation of axle-less TF_1_ at an ATP concentration above *K*_m_. In the expanded trace, the new dwell and catalytic dwell are indicated in green and blue, respectively. The x–y plots and expanded time courses are shown as insets. Videos were recorded at 30,000 fps. (B) Histogram of the dwell time for the new dwell, yielding 𝜏*_N_* = 4.7 ms (N = 1253, 3 molecules). (C) Histogram of the dwell time for the catalytic dwell, yielding 𝜏*_C_* = 3.9 ms (N = 1100, 3 molecules).

The histograms of dwell time for the new dwell and the catalytic dwell were each well fitted by a single exponential decay function, yielding time constants 𝜏*_c_* = 4.7 ms and 𝜏_$_ = 3.9 ms, respectively (Fig. 5B, C). The corresponding exit rates were defined as the inverse of these time constants.

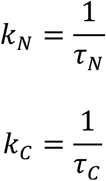

From the catalytic dwell, we counted 80°-forward steps to the next new dwell (𝑛*_f_* = 154) and 40°-backward steps to the new dwell (𝑛*_b_* = 717), giving the forward-branching probability from the catalytic dwell.

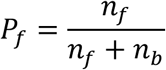

Based on these values, the backward rate constant from the catalytic dwell was defined.

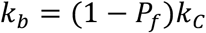

The free-energy bias between the new dwell and the catalytic dwell along the 40° transition was then estimated from these rate constants.

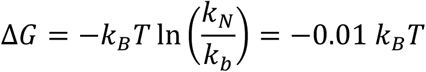

Thus, the new dwell and catalytic dwell states are nearly isoenergetic, consistent with frequent backward steps driven by thermal fluctuations.

### Structural analysis

The structure of axle-less TF_1_ in the presence of ATP was determined by cryo-EM, following a previous study. Axle-less TF_1_ has previously been analyzed by cryo-EM in the presence of AMPPNP as a substrate analogue, and a single state corresponding to the binding dwell was resolved (18). Considering that AMPPNP is a non-hydrolyzable ATP analog, the obtained conformation may not necessarily represent active rotational intermediates under ATP turnover conditions (28). Moreover, among the three distinct dwells indicated by our single-molecule rotation data, two corresponding structural states were not resolved under those conditions in the previous study. To capture conformations that more closely reflect rotary intermediates during ATP-driven turnover, we re-examined the structure of axle-less TF_1_ under ATP substrate conditions.

Axle-less TF_1_ samples purified as described previously (28) were applied to EM grids in presence of ATP at 22°C, vitrified in liquid ethane, imaged at 300 kV, and processed by single-particle analysis (SPA). Under these conditions, we obtained three cryo-EM maps, as expected (Fig. S7). The map resolutions, estimated by the gold-standard procedure, were 2.6 Å (binding dwell), 2.6 Å (catalytic dwell), and 2.5 Å (new dwell) (Fig. S8). However, these maps were highly anisotropic, with 3D estimates suggesting resolutions around the 3 Å range. Overall, the maps showed features that could define the positions and conformations of each subunit (Fig. 6A) and hypothesize the likely nucleotide occupancy (Fig. S11). Given the anisotropic nature of these maps, only the cryo-EM maps were submitted to the EMDB and the models used to interpret the maps are included with this manuscript as supplementary data.

**Fig. 6.**
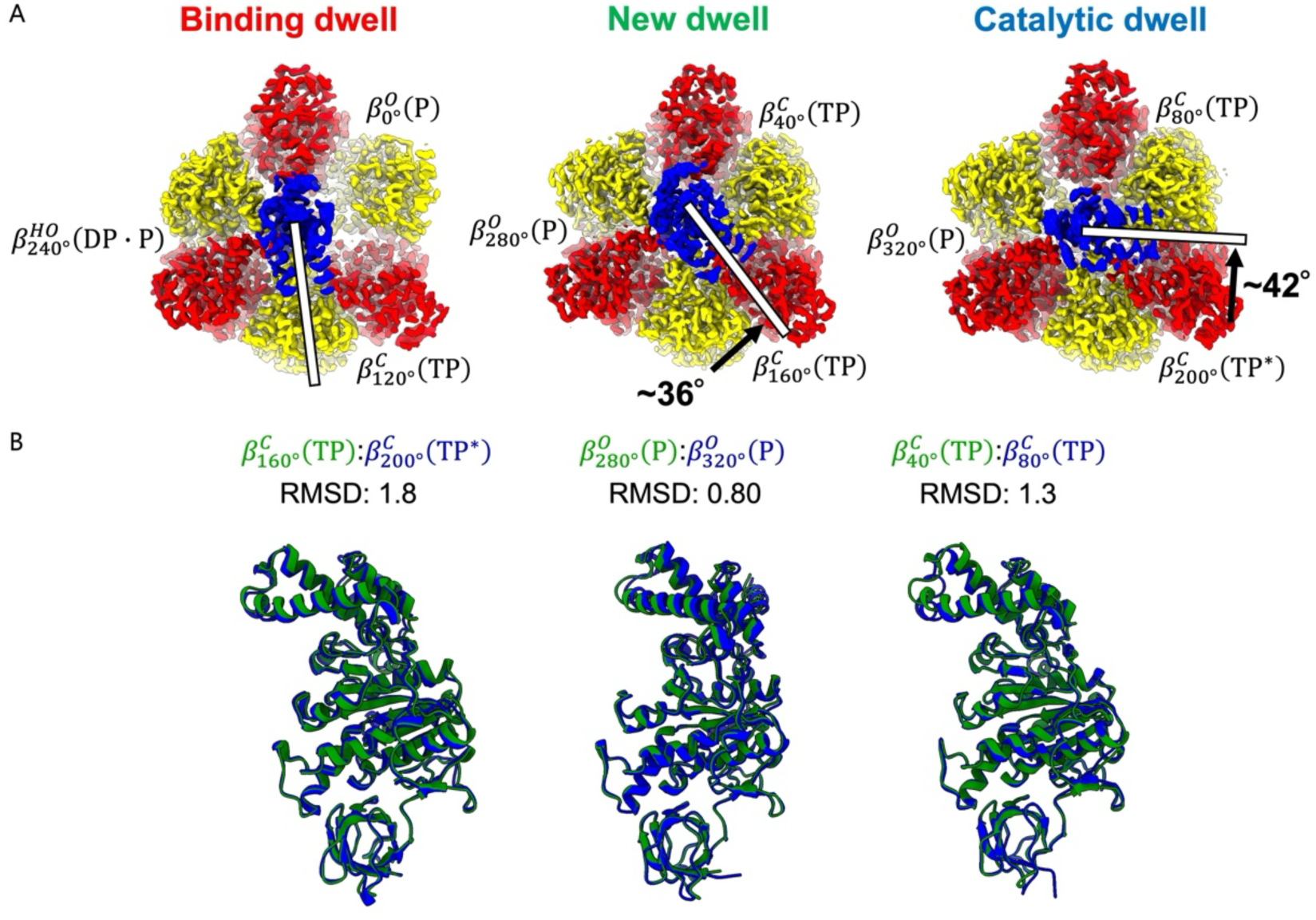
Structure of axle-less TF_1_. (A) Structures of the three rotational dwells of axle-less TF_1_. From left to right: binding dwell, new dwell, and catalytic dwell. Top views are shown, and subunits are colored as in Fig. 1. These structures indicate that the γ subunit rotates counterclockwise among the three dwells (highlighted by a white bar and black arrows). Each dwell exhibits three distinct conformations of the β subunits, representing six substates that the enzyme proceeds through during the hydrolysis cycle 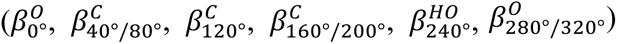. (B) Comparison of β conformations in axle-less TF_1_. The β subunit structures of axle-less TF_1_ new dwell were aligned with the corresponding β structures of axle-less TF_1_ catalytic dwell, with a focus on the N-terminal ∼81 residues that form the β-barrel. The new and catalytic dwell structures are shown in green and blue, respectively.

By comparison with previously reported cryo-EM structures of TF_1_ (PDB IDs: 7L1Q for the binding dwell and 7L1R for the catalytic dwell) (18), we assigned two of the three maps to the binding dwell and catalytic dwell states, respectively. The remaining map exhibited a new structure in which the γ subunit is orientated at an angle intermediate between the binding and catalytic dwell states (Fig. 6A); at +36° from the binding dwell structure. This is in fine agreement with the single-molecule rotation assay showing the new dwell state at +41° from the binding dwell. The angular difference between the new dwell and catalytic dwell states was 42°, also consistent with the 39° difference determined by the rotation assays (Fig. 6A). Thus, the new structure was interpreted as the new dwell state identified by the single-molecule rotation assays.

Structural comparison of the β subunits between axle-less TF_1_ and WT TF_1_ revealed highly similarity (Fig. S9). In addition, the β conformations in the binding dwell state closely matched those in the previously reported axle-less TF_1_ binding dwell structure (Fig. S10; (28). RMSD values (Å) represent the structural differences between the aligned models.

The cryo-EM density clearly indicates whether a nucleotide is present at each catalytic site (Fig. S11). However, ATP vs. ADP identity and Pi-bound vs. empty assignments are interpretations of the cryo-EM density, and should be treated as such rather than definitive conclusions.

Comparison of the new dwell and catalytic dwell structures of axle-less TF_1_ showed that these two states primarily differ in the rotational angle of the γ subunit, whereas the β subunit conformations remain closely aligned (Fig. 6B). This structural similarity suggests that a large β conformational power stroke is unlikely to drive the transition between these two states. Instead, the 40° rotation appear to be thermally driven, consistent with the backstep analysis.

As with nucleotide assignments, the cryo-EM maps did not contain sufficient detail to assign all residue rotamers de novo. However, residues positions and likely interactions could be inferred from the rotational position of γ and the absence of detectable conformational changes within the subunits relative to previously published structures.

Using these models and maps, we compared interactions between the α_3_β_3_ ring and γ in the new dwell and the catalytic dwell. This comparison suggested that the new dwell may contain novel interactions that are absent in the catalytic dwell. When α and β residues located within 4.5 Å of γ were identified, 𝛽_280°_E379–γM29, αE391–γQ22, αA394–γK19, αA394–γI23, and αF398–γI23 were detected exclusively in the new dwell (Fig. S15). All the α residues refer to the α subunit positioned between 𝛽_280°_ and 𝛽_160°_. Among these, 𝛽_280°_E379–γM29 and αA394–γK19 are likely to form van der Waals interactions, αE391–γQ22 a hydrogen bond, and αA394–γI23 and αF398–γI23 hydrophobic interactions. Taken together, these findings suggest that these interactions contribute to stabilization of the rotational pause at the new dwell.

Another aspect found in the new structures of axle-less TF_1_ is the tilt of the γ subunit. Whereas obvious off-axis tilt was not found in the binding dwell structure, the catalytic dwell structure exhibits a noticeable tilt, when compared with the corresponding structures of WT TF_1_. The tilt angles between axle-less TF_1_ and WT TF_1_ were quantified as 1.3° and 7.5° for the binding dwell and catalytic dwell states, respectively. To test whether the tilt can be accommodated only in axle-less TF_1_, we generated models in which the axle-less γ was replaced with the full-length γ. Aligning the γ subunit of WT TF_1_ in the catalytic dwell state to the γ subunit in axle-less TF_1_ resulted in steric clashes with the lower region of α and β subunits (Fig. S12). These results suggest that the off-axis tilt is enable by the loss of the lower γ axle and help avoid steric interference during rotation.

### Rotational scheme model of axle-less TF_1_

Combining the results from single-molecule rotation assays and cryo-EM structural analysis, we propose a rotational scheme model of axle-less TF_1_ (Fig. 7; Movie S1). The angular positions of the binding and catalytic dwell matched those of WT TF_1_. In addition, axle-less TF_1_ exhibited a unique rotational dwell state—termed the “new dwell”—located ∼40° between the binding and catalytic dwell. In the new dwell state, the conformation of the catalytic β subunits closely matches that in the catalytic dwell states 40° ahead. Moreover, the free-energy difference between the new dwell and catalytic dwell is less than 1𝑘*_B_*𝑇, suggesting that the structural transition between these intermediates can occur within the range of thermal fluctuations.

**Fig. 7.**
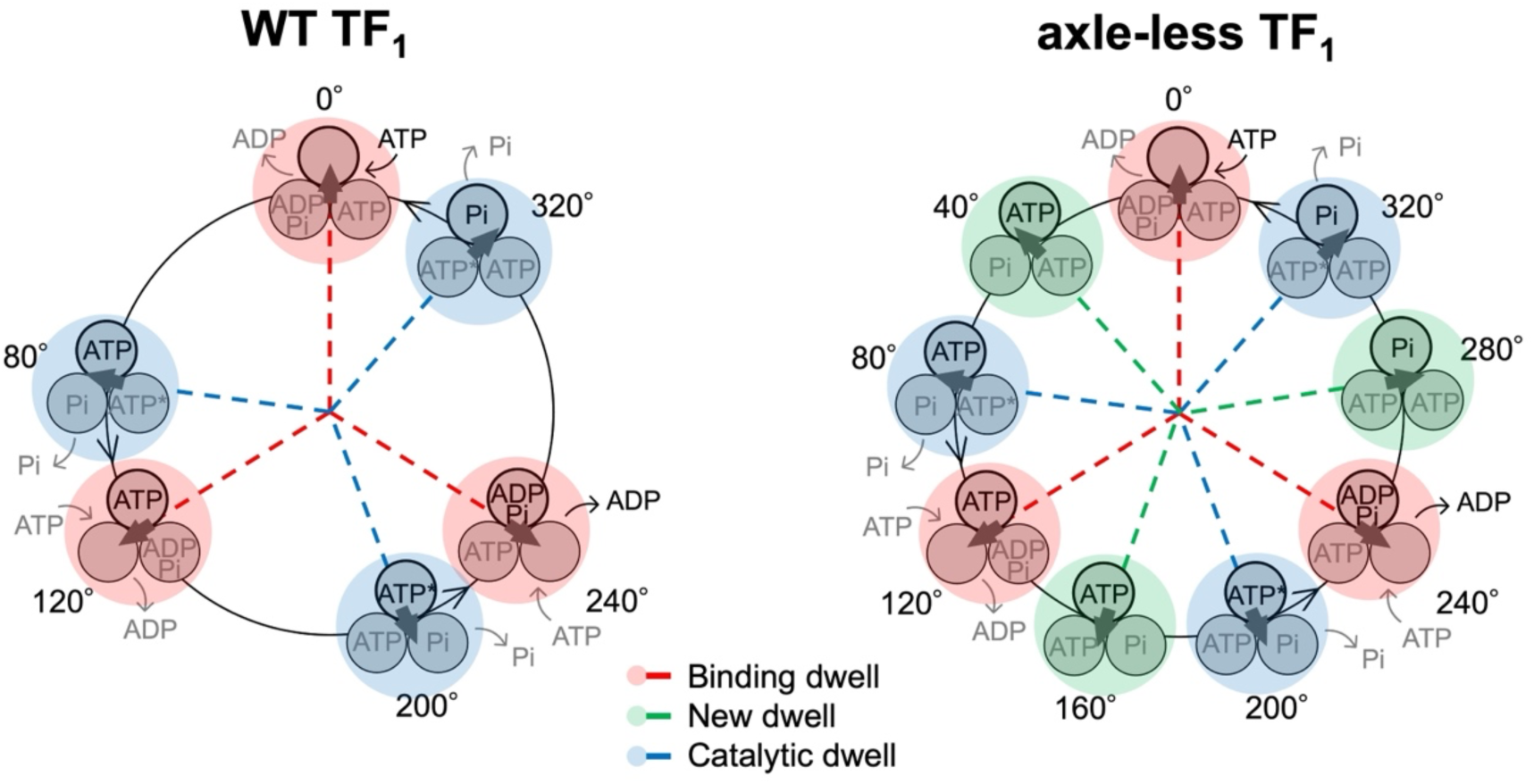
Rotational scheme model of WT TF_1_ and axle-less TF_1_. The left panel shows WT TF_1_ and the right panel shows axle-less TF_1_. The three circles represent the three β subunits in the stator ring, and the nucleotides predicted to occupy the binding sites of each β subunit are shown inside the circles. The arrow indicates the rotational angular position of the γ subunit. The 0° position is defined as the γ angle at which a given β subunit binds ATP. An asterisk following “ATP” denotes the catalytically competent state in which the bound ATP undergoes hydrolysis. The angular positions of the binding dwell, catalytic dwell, and new dwell are indicated in red, blue, and green, respectively.

## Discussion

We investigated the functional role of the lower region of the γ axle by analyzing axle-less TF_1_, a mutant lacking the lower half of the γ axle. By combining single-molecule rotation assays with cryo-EM structural analysis, we identified a previously unrecognized intermediate state, termed the “new dwell,” located approximately midway between the binding dwell and the catalytic dwell. High-time-resolution single-molecule measurements further revealed that the transition between the new dwell and the catalytic dwell is reversible, with frequent backward transitions from the catalytic dwell to the new dwell.

Consistent with these observations, cryo-EM analysis showed that the new dwell and catalytic dwell share essentially the same α_3_β_3_ stator-ring conformation, with two β subunits in the closed conformation and one β subunit in the open conformation (the CCO state). These findings indicate that the axle-less γ subunit can adopt two distinct stable angular positions within the same stator-ring conformation. The comparable particle populations of the two states further suggest that they are nearly isoenergetic. In contrast, the binding dwell and new dwell structures exhibited clear differences in β-subunit conformations. Taken together, these results suggest that the lower region of the γ axle plays a critical role in maintaining unidirectional rotation and torque generation specifically during the latter half of the 0–80° rotational substep.

Based on these findings, we propose the reaction scheme for axle-less TF_1_. Although the current cryo-EM density around the catalytic site does not allow definitive assignment of the nucleotide chemical states, our results suggest that ATP cleavage occurs not at the new dwell, but at the catalytic dwell located ∼80° ahead of the binding dwell. Under ATPγS conditions, the catalytic pause was markedly prolonged in the single-molecule rotation assay. In addition, high-frame-rate recordings showed that forward rotation proceeded after the pause at the catalytic dwell position. Together, these observations support assignment of ATP cleavage to the 80° position. Therefore, despite the overall structural similarity between the new dwell and catalytic dwell states, some differences must remain in the catalytic sites. However, the present density maps do not yet allow reliable evaluation of subtle catalytic features, including the configuration of residues directly involved in hydrolysis such as the arginine finger (6).

In a previous structural study of axle-less TF_1_ (28), only the binding dwell state was resolved, whereas structures corresponding to the new dwell and catalytic dwell were not observed. This discrepancy may be attributable to the use of AMPPNP, a non-hydrolyzable ATP analog. AMPPNP is commonly used to arrest F_1_ rotation at the catalytic dwell, which is thought to correspond to a pre-hydrolysis state. However, the previous study reported a binding-dwell structure together with catalytic sites occupied by nucleotide hydrolysis products. These observations raise the possibility that AMPPNP did not fully inhibit rotation of axle-less TF_1_ and that residual ATP contamination instead drove the motor under ATP-binding-limited conditions. Consistent with this interpretation, our single-molecule experiments at ATP concentrations far below *K*_m_ detected only three pauses corresponding to the binding dwell, which may explain why the new dwell and catalytic dwell structures were not captured in the previous cryo-EM analysis.

One additional structural implication emerged from the present study. Because the new dwell structure has not been identified in previous analyses of WT TF_1_ containing the full-length γ subunit, the new dwell is likely to be specific to axle-less TF_1_. Indeed, when the full-length γ subunit from WT TF_1_ was inserted into the new dwell structure, severe steric clashes with the stator ring were observed, supporting this interpretation (Fig. S13). However, the γ conformations in the binding dwell and catalytic dwell are not identical. We therefore generated an intermediate γ model between these two conformations and inserted it into the stator ring of the new dwell structure at the +40° angular position. Interestingly, in this model, steric clashes were largely avoided even without introducing the γ tilt observed in axle-less TF_1_ (Fig. S14). This result suggests that a γ subunit positioned at +40° relative to the CCO-state stator ring may in principle exist transiently in WT TF_1_. It may therefore be necessary to examine whether such a state appears during the rotary cycle of WT TF_1_. At the same time, however, γ rotation without accompanying conformational changes in the catalytic β subunits appears inconsistent with early theoretical studies (29), our previous single-molecule manipulation experiments (30), and recent molecular dynamics simulations (31), all of which support tight coupling between γ rotation and β conformational transitions.

## Methods

### Preparation of F_1_

WT TF_1_ for single-molecule rotation assays was prepared as described previously (32). To attach 40-nm gold colloids for rotation observation, two cysteine residues were introduced (33) and the F_1_ complex was biotinylated (11).

The plasmid encoding axle-less TF_1_ was constructed using the TF_1_ plasmid pHCXP as the vector and the plasmid encoding γΔN4C25 (pHC95_γΔN4C25; Furuike et al., 2008) as the insert. After ligation, the recombinant plasmid was transformed into the F_o_F_1_-deficient *E. coli* strain JM103Δunc. Axle-less TF_1_ was expressed in *E. coli* and purified and biotinylated using the same procedure as for WT TF_1_. Protein concentration was determined from the absorbance at 280 nm using a molar extinction coefficient of 154,000.

For cryo-EM structural analysis, axle-less TF_1_ protein (γΔN4C25) that was purified in (28) was used. The protein was snap frozen and stored at -80℃ until utilized in the present study.

### Single-molecule rotation assays

To observe F_1_ rotation, 40-nm gold colloids (prepared as described in Iida et al., 2019) were attached to the biotinylated γ subunit as optical probes.

Flow cells were constructed from two cover glasses (18 × 24 mm^2^ and 24 × 32 mm^2^; Matsunami Glass) using double-sided tape as a spacer. To capture and immobilize F_1_ by His-tags introduced into the α_3_β_3_, the bottom glass surface was coated with Ni-NTA.

The single-molecule rotation buffer contained 50 mM MOPS-KOH (pH 7.0), 50 mM KCl, and 2 mM MgCl_2_. When ATP was used as the substrate, an ATP-regenerating system (100 µg/mL pyruvate kinase and 2.5 mM phosphoenolpyruvate) was included.

For the rotation assay, the flow cell was first filled with rotation buffer containing 5 mg/mL BSA (BSA buffer) and incubated for 5 min. F_1_ in the BSA buffer (WT TF_1_, 200 pM; axle-less TF_1_, 3 nM) was then introduced and incubated for 5 min, followed by washing with BSA buffer to remove unbound molecules. Next, 40-nm gold colloids were introduced and incubated for 10 min. Finally, the flow cell was washed with rotation buffer containing substrate to remove unbound particles.

Rotation assays were performed at room temperature as described in Iida et al., 2019. Rotation was observed with a dark-field microscopy (IX-71, OLYMPUS) with a 60× objective lens and recorded using a high-speed camera (FASTCAM-1024PCI, Photron) at 500–30,000 fps.

### Single-molecule data analysis

Recorded videos were analyzed using custom software. Pause angles were determined by fitting the angular position histograms with Gaussian functions (Fig. 3B and 4B). For backward step analysis, change point analysis was applied to the time traces (Fig. 5A) to automate pause detection and averaging, as described previously (35, 36). To estimate time constants, the histograms of dwell time were fitted with single exponential decay functions.

### Cryo-EM grid preparation

Purified axle-less TF_1_ (3.5 µL) was supplemented with 20 mM ATP and MgCl_2_ and applied to a glow-discharged holey gold grid (UltrAfoil R 1.2/1.3, 200 mesh). The grid was blotted for 4 s at 22℃ with a blot force of 0 and a chamber humidity of 100%, and plunge-frozen in liquid ethane using an FEI Vitrobot Mark IV. The time from ATP addition to freezing was ∼15 seconds.

### Data collection

Grids were transferred to a Thermo Fisher Talos Arctica transmission electron microscope (TEM) operating at 200 kV and screened for ice thickness and particle density. Grids were subsequently transferred to a Thermo Fisher Titan Krios TEM G3i operating at 300 kV equipped with a Gatan BioQuantum energy filter (15 eV slit) and K3 Camera at Molecular Horizons, University of Wollongong. Images were recorded automatically using EPU at ×59,000 (displayed magnification of ×105,000 due to the energy filter), resulting in a pixel size of 0.84 Å. A total dose of 74 electrons per Å^2^ was used spread over 80 frames, with a total exposure time of 8.1 s. 13,845 movie micrographs were collected.

### Cryo-EM data analysis

All image processing and refinements were performed in cryoSPARC v4.4.1 (37) using default settings unless otherwise specified. Raw movies were first motion-corrected, and CTF parameters were estimated using the patch-based workflow with default parameters (maximum alignment resolution, 5 Å; alignment B-factor, 500; defocus search range, 1,000–40,000 Å). Micrographs with relative ice thickness greater than 1.05 or estimated CTF resolution worse than 4.0 Å were removed to yield a final set of 11,193 movies micrographs. Particles were initially auto-picked on a subset of the dataset (200 micrographs) using Blob Picker (particle diameter, 100–150 Å) to generate templates for picking. Then, using Template Picker, a total of 7,720,880 particles were picked from the full dataset of 11,193 micrographs. Picked particles were extracted with a box size of 256 pixels, and multiple rounds of 2D classification were performed to exclude “junk” particles derived from aggregates or minor contaminants at the end of each round. After 2D classification, 3,736,633 particles that appeared to represent the complex were retained.

Ab initio and heterogeneous refinement produced three distinct classes: a class corresponding to the binding dwell (608,830 particles), a class corresponding to the catalytic dwell (845,464 particles), and a class corresponding to the new dwell (852,517 particles). Each of the three maps was further subjected to 3D classification using a mask on the rotor γ subunit, and classes with clear γ density were selected. The selected particle numbers were 422,075 for the binding dwell, 415,389 for the catalytic dwell, and 464,363 for the new dwell. Each map was then refined independently by non-uniform refinement to obtain the final high-resolution maps. Particle numbers at each step are detailed in the supplementary flowchart (Fig. S7), and statistics for all final maps and models are listed in Fig. S8 and Table S1. Because some regions exhibited low-resolution features, DeepEMhancer (38) was applied to sharpen the maps and improve interpretability for figures showing the entire complex. Non-uniform refined maps were uploaded to the 3DFSC processing server to obtain additional 3D FSC information.

### Model building

Atomic models were built for the intact F_1_ complex and refined using Coot (39), Isolde (40) and PHENIX (41), with the TF_1_ cryo-EM structures 7L1Q (binding dwell) and 7L1R (catalytic dwell) serving as templates. Due to the anisotropic nature of the maps, pdbs were not submitted, instead they are attached as supplementary. Figures were prepared using ChimeraX-1.10.1.

### Estimation of the γ subunit rotational angle and tilting

The rotational displacement of the γ subunit between the binding and new dwell states and between the new and catalytic dwell states was measured in ChimeraX-1.10.1 using the “Measure Rotation” command. The axis of the central γ helix was defined by the Cα atoms of residues 12-42 and 221-257 in chain G. Using this axis definition, the angular differences were 36° (binding to new dwell) and 42° (new to catalytic dwell). Because the γ axis is tilted, these values include an uncertainty of approximately ±5°.

The tilt of the γ subunit was also quantified in ChimeraX-1.10.1. The γ-axis was defined with the “define axis” command using the Cα atoms of residues 10-46 and 223-261 in chain G for the binding dwell state, and residues 12-52 and 220-261 in chain G for the catalytic dwell state. For each dwell, the tilt was defined as the angle between the γ-axis of WT TF_1_ and that of axle-less TF_1_ and was measured with the “angle” command. Using this procedure, the γ tilt was 1.3° in the binding dwell and 7.5° in the catalytic dwell.

### Construction of TF_1_ models with an artificially 40°-rotated γ subunit

Models were generated in which the γ subunit of TF_1_ was artificially rotated by +40° in the binding dwell structure (PDB ID: 7L1Q) and by -40° in the catalytic dwell structure (PDB ID: 7L1R). First, each F_1_ structure was translated so that the center of the three centers of mass (Cα atoms only) of the N-terminal domains (residues 10–82) of the three β subunits became the origin. Next, the structure was rotated so that the z axis became perpendicular to the plane defined by these three centers of mass. The F_1_ complex was then rotated around the z axis so that the center of mass of the N-terminal domain (residues 10–82) of the β subunit (chain D) lay on the x axis, thereby aligning the α_3_β_3_ ring in a common reference frame. After ring fitting, the coordinates of the γ subunit (chain G) were rotated about the z axis passing through the origin by applying a rotation matrix 𝑅_1_(𝜃) only to γ, while keeping the α and β subunits fixed. The rotation angle was set to θ = +40° for the 7L1Q-derived model and θ = -40° for the 7L1R-derived model. The rotated γ coordinates were then merged back with the unmodified ring coordinates to generate TF_1_ models with an artificially rotated γ subunit.

## Supporting information

Binding dwell

New dwell

Catalytic dwell

## Acknowledgements

We thank all the members of our laboratory for their helpful comments and discussions. The author gratefully acknowledges support from MERIT-WINGS. This study was supported in part by a Grant-in-Aid for Scientific Research on Innovative Areas (JP21H00388 to H.U.), a Grant-in-Aid for Challenging Research (Exploratory; JP23K18092 to H.U.), a Grant-in-Aid for Scientific Research (B) (JP24K01987 to H.U.), and a Grant-in-Aid for Scientific Research (S) (JP19H05624 to H.N.) from JSPS, and a Research Grant from Human Frontier Science Program (Ref. No: RGP0054/2020 to H.N.), and a JST ASPIRE Program (JPMJAP24B5 to H.N.). We wish to thank and acknowledge the use of the University of Wollongong Cryogenic Electron Microscopy Facility at Molecular Horizons, as well as the use of the Victor Chang Cardiac Research Institute Innovation Center (funded by the NSW Government) and the Electron Microscope Unit at UNSW Sydney. A.G.S. is supported by National Health and Medical Research Council grants 2016308 and this research was partially funded by the Australian Research Council through Discovery Project DP250101405. This research was conducted by the Australian Research Council Industrial Transformation Training Centre for Cryo-Electron Microscopy of Membrane Proteins for Drug Discovery (IC200100052).

## Supporting information

Additional supporting information is available and is provided as separate files. The molecular models used for interpretation are provided in pdb format.

## Data availability

Cryo-EM maps have been deposited under Electron Microscopy Data Bank (EMDB) codes EMD-69782, EMD-69783 and EMD-69784. Molecular models are included as supplementary data.

