## Supporting Information for "The Lower γ Region Ensures Unidirectional Rotation and Torque Generation in the Latter Half of the 80° Substep of F1-ATPase"

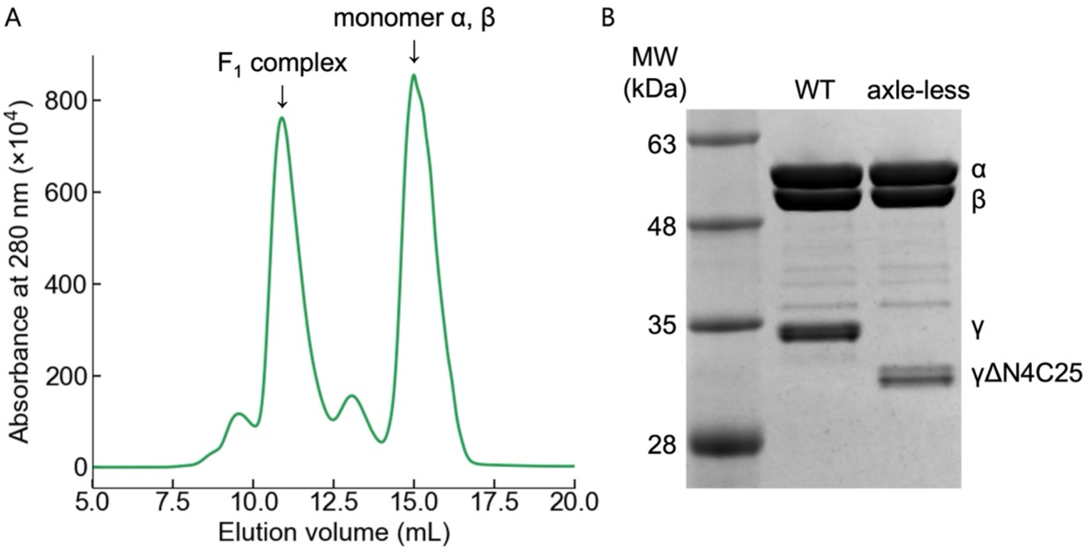


Fig. S1. Gel filtration chromatogram and SDS-PAGE analysis of axle-less $TF_1$.

(A) Gel filtration chromatogram of axle-less $TF_1$ after Ni-NTA purification. The peak corresponding to the $F_1$ complex eluted at nearly the same position as the peak observed during purification of WT $TF_1$ with a comparable molecular weight. Peaks corresponding to α and β monomers were also detected. (B) SDS-PAGE analysis of axle-less $TF_1$. From left to right, lanes correspond to the marker, WT $TF_1$, and axle-less $TF_1$. WT $TF_1$ and axle-less $TF_1$ were loaded at the same concentration. The γ subunit of axle-less $TF_1$ was detected at a lower apparent molecular weight than that of WT $TF_1$. The α and β band intensities were comparable between WT $TF_1$ and axle-less $TF_1$, whereas the γ band was weaker in axle-less $TF_1$, consistent with reduced γ content in the axle-less $TF_1$. The γ subunit showed doublet bands due to partial biotinylation, as reported previously (1).

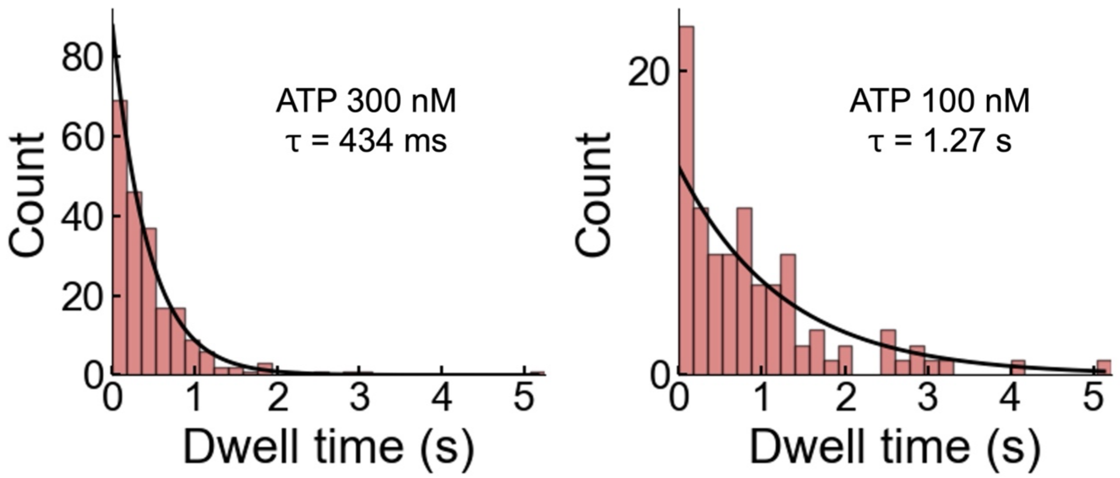


Fig. S2. Time-constant analysis at low [ATP] far below $K_m$

Histograms of the dwell time at 300 nM ATP (N = 214, 4 molecules) and 100 nM ATP (N = 99, 4 molecules). Videos were recorded at 500 fps.

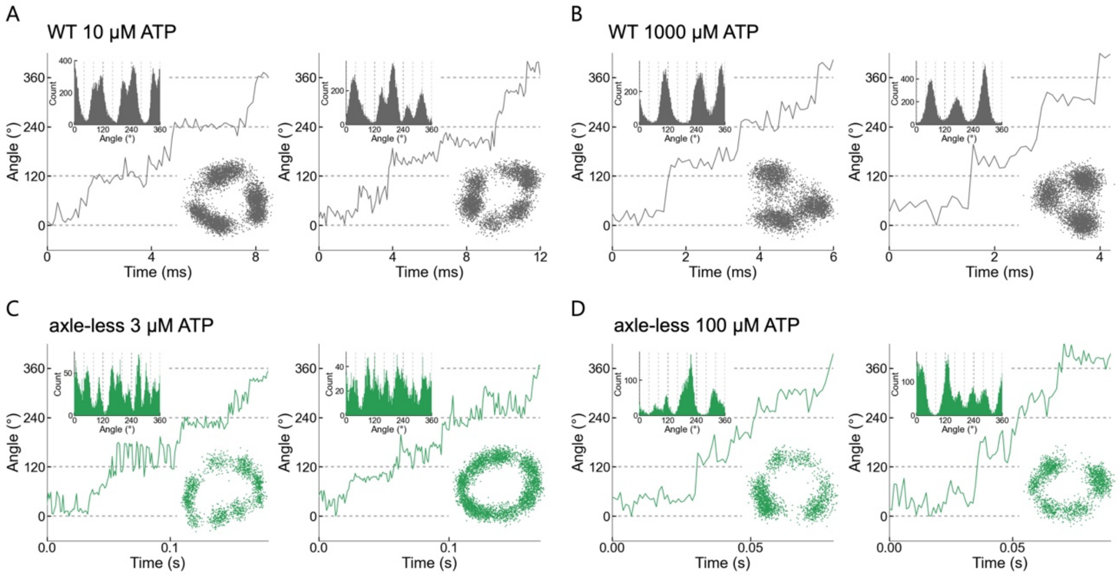


Fig. S3. Single-molecule rotation assays of WT $TF_1$ and axle-less $TF_1$, related to Fig. 2.

Representative rotation time courses. The x–y plots (right) and angular position histograms (left) are shown as insets. Two molecules are shown as examples for each condition. (A, B) Rotation of WT $TF_1$ recorded at 10,000 fps. (A) 10 μM ATP. (B) 1000 μM ATP. (C, D) Rotation of axle-less $TF_1$ recorded at 1,000 fps. (C) 3 μM ATP. (D) 100 μM ATP.

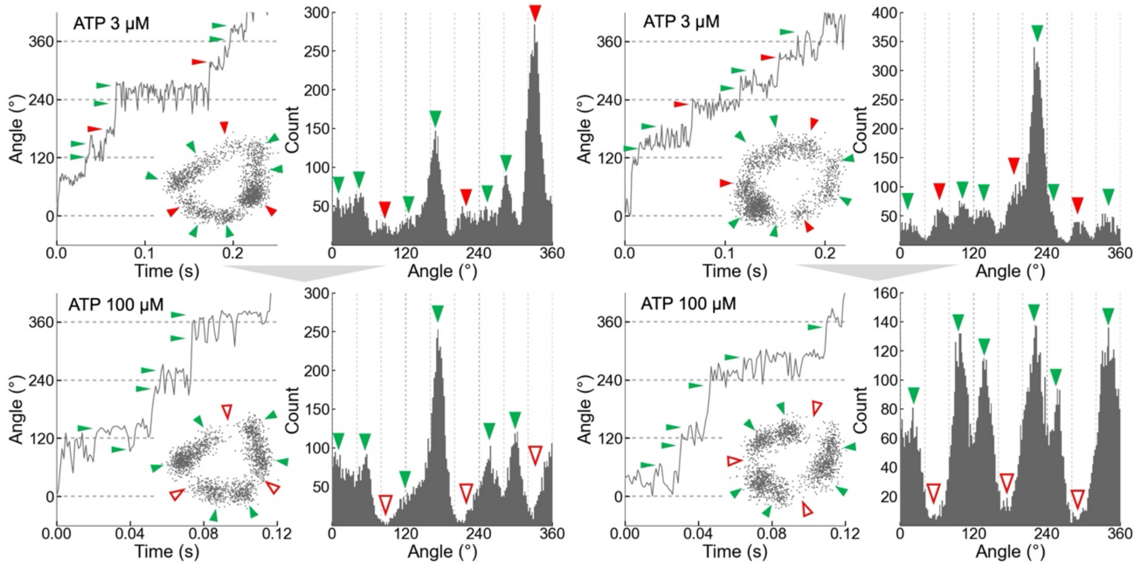


Fig. S4. Identification of the binding dwell from buffer exchange experiments, related to Fig. 3.

Rotation time courses and angular histograms for the same molecule during a buffer exchange experiment, in which [ATP] was switched between a low concentration around the $K_m$ range (top) and a high concentration above $K_m$ (bottom). The x–y plots are shown as insets. Two molecules are shown as examples. Pauses indicated by red arrows were identified as binding dwells. Among the pauses indicated by green arrows, three are inferred to correspond to catalytic dwells and the remaining three to an additional dwell state not observed in WT $TF_1$. Videos were recorded at 1,000 fps.

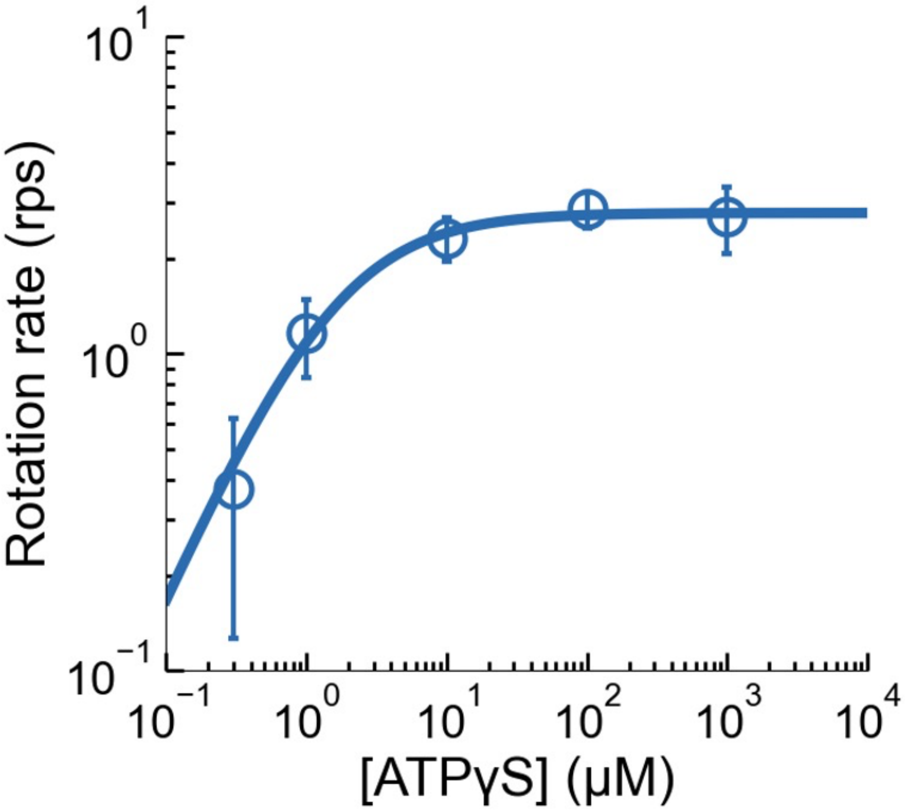


Fig. S5. [ATPγS] versus rotation rate.

[ATPγS] is plotted against the rotation rate for axle-less $TF_1$. The mean and SD of each data point are shown as circles and error bars, respectively (n = 4). Solid lines indicate Michaelis–Menten fits. $V_{max}$: 2.8 rps, $K_m$: 1.6 μM. The ATPγS-binding rate constant ($k_{on}$), estimated as $3 \times V_{max}/K_m$, was $5.3 \times 10^7$ $M^{-1}$ $s^{-1}$.

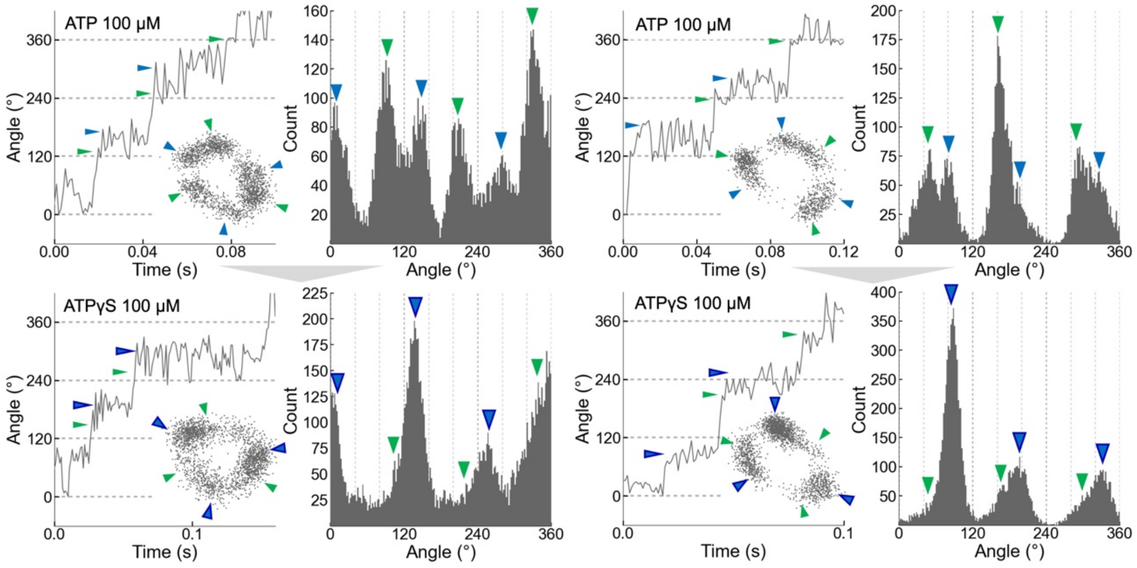


Fig. S6. Identification of the catalytic dwell from buffer exchange experiments, related to Fig. 4.

Rotation time courses and angular histograms for the same molecule during a buffer exchange experiment, in which the substrate was switched between ATP (top) and ATPγS (bottom) at concentrations above $K_m$. The x–y plots are shown as insets. Two molecules are shown as examples. Pauses indicated by blue arrows were identified as catalytic dwells. Pauses indicated by green arrows were termed as the “new dwell,” unique to axle-less $TF_1$. Videos were recorded at 1,000 fps.

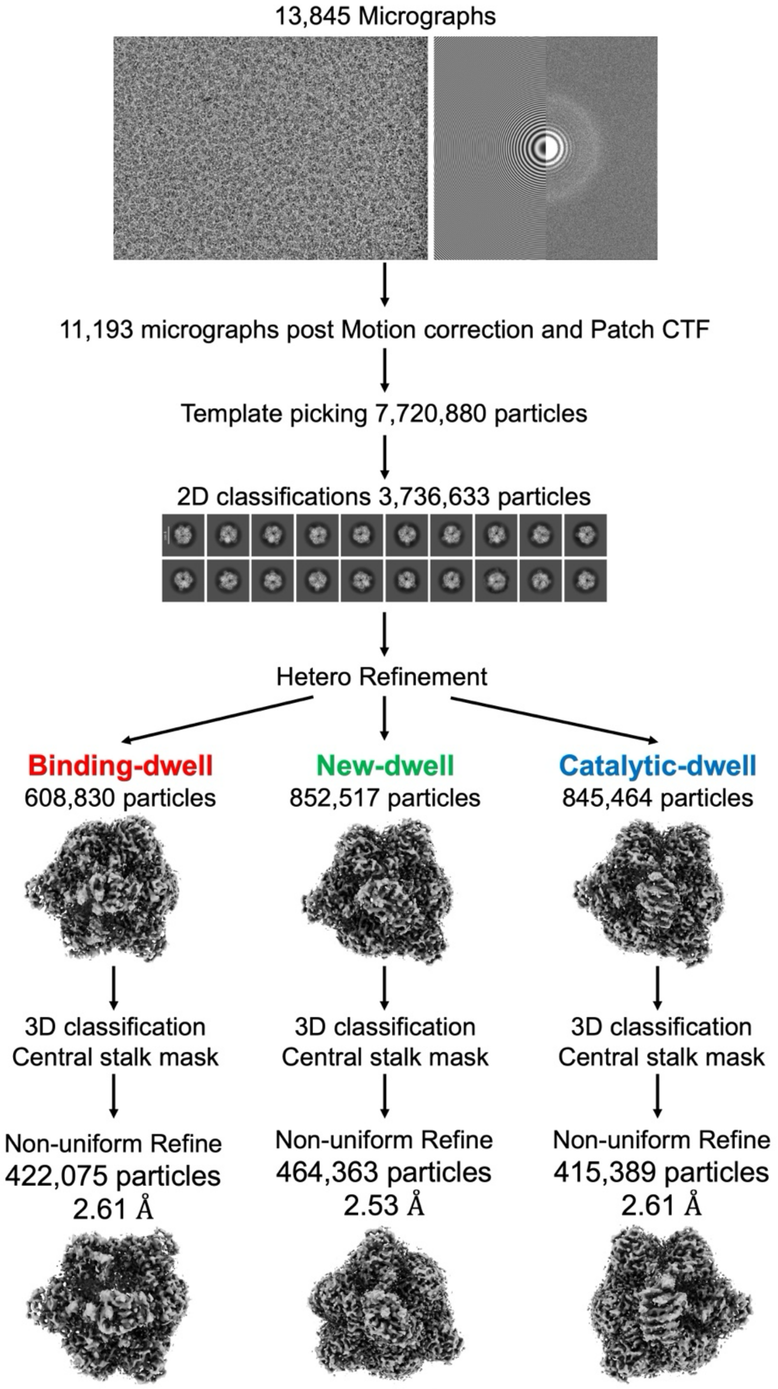


Fig. S7. Cryo-EM data collection and data processing flowchart.

The white scale bar in the 2D class-average images corresponds to 100 Å. The dataset was separated into substates using heterogeneous refinement.

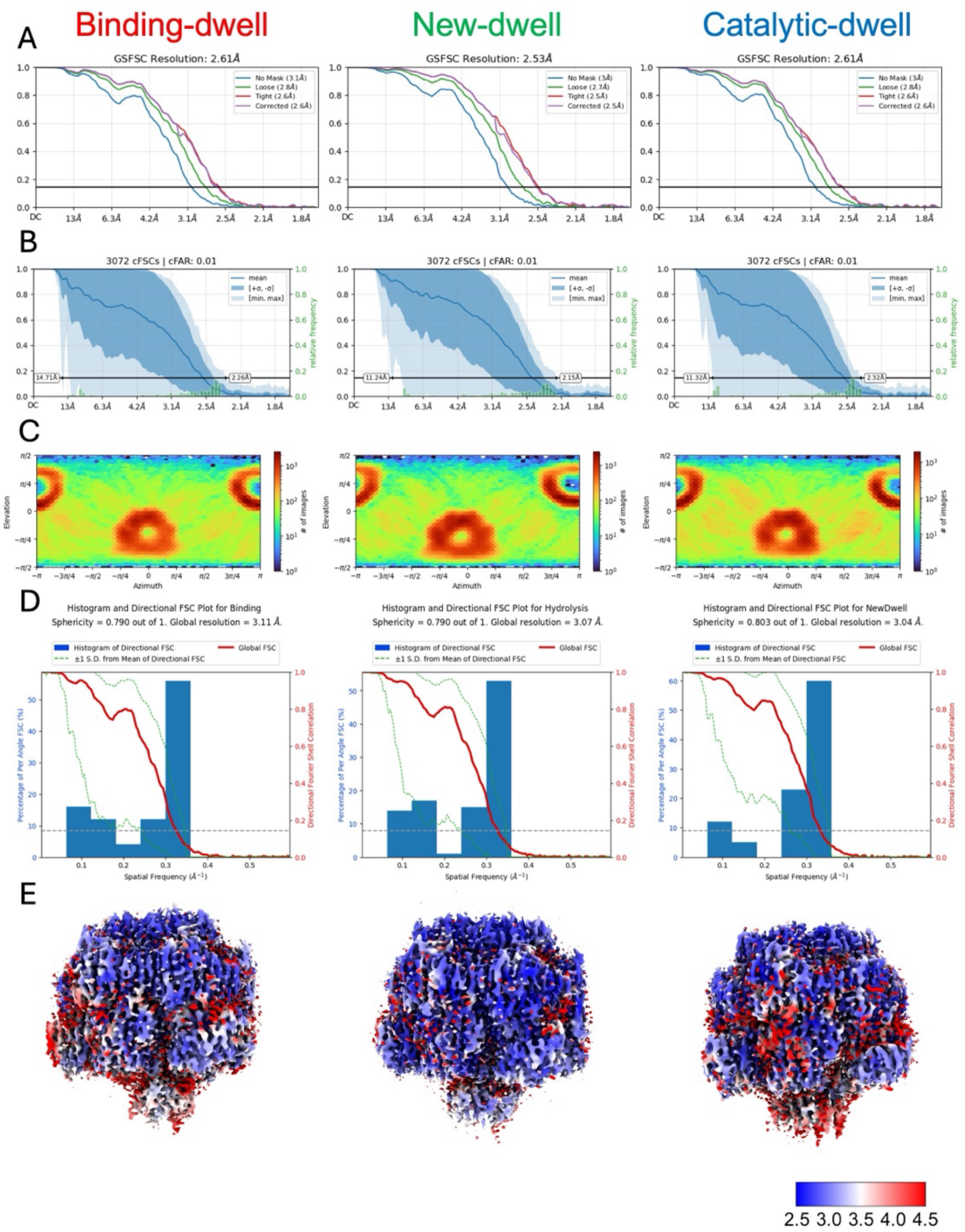


Fig. S8. 3D FSC Curves and Local Resolution Estimates.

(A) Gold-standard Fourier shell correlation (GSFSC) curves calculated in cryoSPARC. (B) Conditional Fourier shell correlation (cFSCs) curves calculated in cryoSPARC. (C) Viewing direction distribution plot. (D) Histograms and direct FSC plots from 3D FSC analysis. (E) Local-resolution estimates calculated in cryoSPARC. The lower-right panel shows the local-resolution scale.

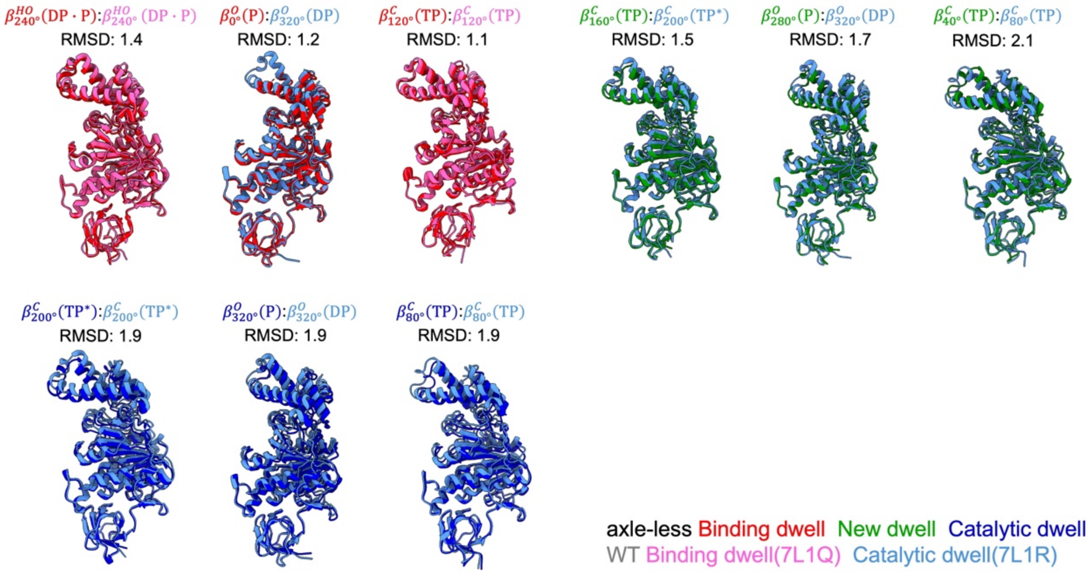


Fig. S9. Comparison of β conformations in axle-less $TF_1$.

The β subunit structures of axle-less $TF_1$ were aligned with the corresponding β structures of $TF_1$, with a focus on the N-terminal ~81 residues that form the β-barrel. The axle-less $TF_1$ binding dwell, new dwell, and catalytic dwell structures are shown in red, green, and blue, respectively, whereas the $TF_1$ binding dwell (PDB ID: 7L1Q) and catalytic dwell (PDB ID: 7L1R) structures are shown in pink and light blue, respectively. Note that $\beta_{0^\circ}$ in the axle-less $TF_1$ binding dwell structure is closer to $\beta^{O}_{320^\circ}$ in 7L1R rather than to the corresponding $\beta^{HC}_{0^\circ}$ in 7L1Q. This feature was also observed in the ATP-waiting state of $TF_oF_1$ (2). This is likely because the binding dwell structure obtained here reflects the ATP-waiting state itself rather than the binding dwell structure that reflects the temperature sensitive (TS) reaction.

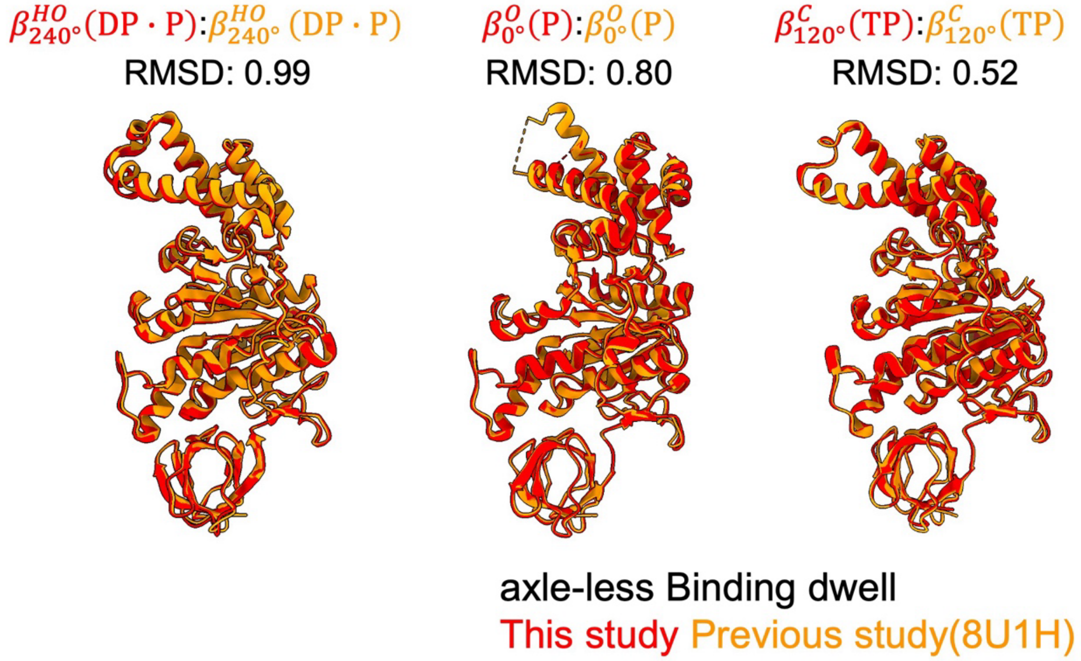


Fig. S10. Comparison of binding dwell β conformations between the axle-less $TF_1$ structure determined in this study and the previous study

The binding dwell β subunit structures of axle-less $TF_1$ of this study were aligned with the corresponding β structures of previously reported axle-less $TF_1$, with a focus on the N-terminal ~81 residues that form the β-barrel. The axle-less $TF_1$ binding dwell structure determined in this study and the previous study are shown in red and orange, respectively. The two structures were in good agreement.

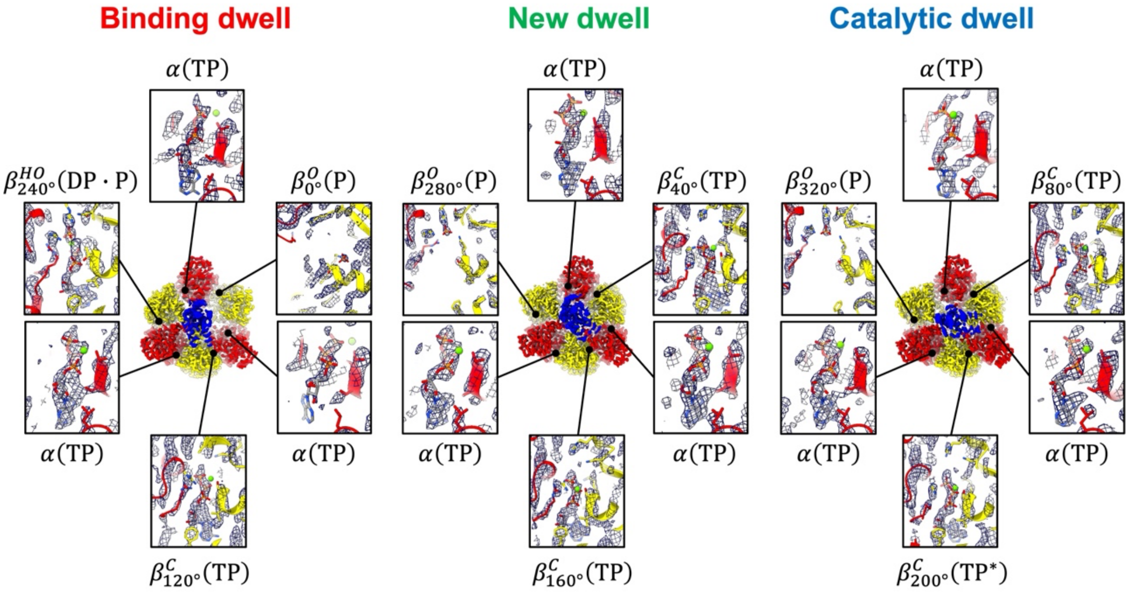


Fig. S11. Close-up views of the nucleotide binding sites.

Cryo-EM maps are shown together with close-up views of each nucleotide binding site. All α subunits in all dwells bound nucleotides. In the binding dwell, $\beta_{0°}$ was nucleotide-free, whereas the equivalent site in $TF_1$ was ATP-bound (3). This difference is likely because the binding dwell structure captured here reflects an ATP-waiting state rather than a binding dwell state that reflects the TS reaction. In the catalytic dwell, $\beta_{320°}$ was nucleotide-free, whereas the equivalent site in $TF_1$ was ADP-bound (3). This difference is likely because the ADP rebinding inferred in $TF_1$ did not occur in axle-less $TF_1$. The nucleotide state in the new dwell closely matched that in the catalytic dwell.

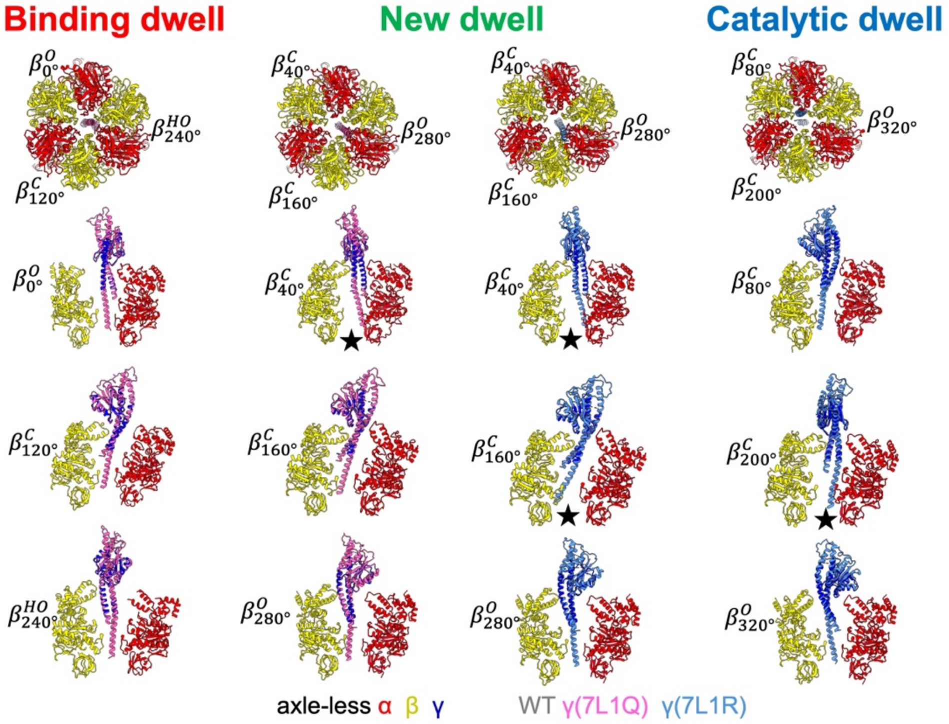


Fig. S12. Steric clashes between γ and αβ upon alignment of $TF_1$ γ onto axle-less $TF_1$ γ.

The $TF_1$ γ subunit structures were aligned with the corresponding γ structures of axle-less $TF_1$, focusing on the N-terminal residues 12–45 and the C-terminal residues 221–257 that form the axial helix. In the representation, the α, β, and γ subunits of axle-less $TF_1$ are shown in red, yellow, and blue, respectively. The γ subunit of $TF_1$ in the binding dwell (PDB ID: 7L1Q) is shown in pink, and the γ subunit of $TF_1$ in the catalytic dwell (PDB ID: 7L1R) is shown in light blue. In the axle-less $TF_1$ binding dwell, the aligned $TF_1$ γ did not show steric clashes with α and β. In the catalytic dwell, the aligned $TF_1$ γ clashed with the α subunit facing $\beta_{200^\circ}$. In the new dwell, the aligned $TF_1$ γ from the $TF_1$ binding dwell clashed with the α subunit facing $\beta_{40^\circ}$, and the aligned $TF_1$ γ from the $TF_1$ catalytic dwell clashed with the α subunit facing $\beta_{40^\circ}$ and with $\beta_{160^\circ}$. Sites of steric clash are indicated by black stars.

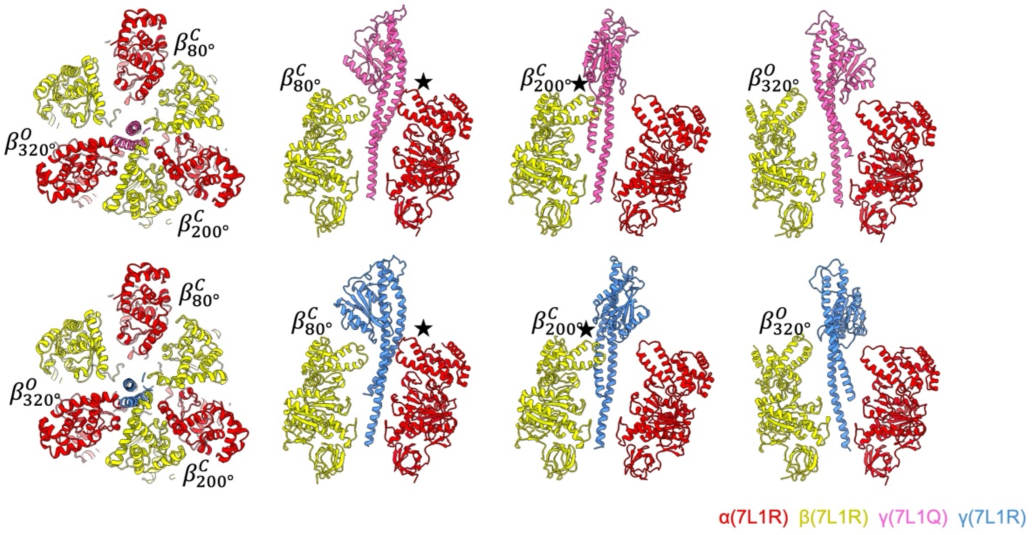


Fig. S13. Steric clash between the $TF_1$ catalytic dwell $\alpha_3\beta_3$ ring and γ subunits at 40° rotational angle

The $TF_1$ γ subunit was rotated relative to the $\alpha_3\beta_3$ ring by +40° for the binding dwell structure and by −40° for the catalytic dwell structure. In the representation, the α and β of $TF_1$ in the catalytic dwell (PDB ID: 7L1R) are shown in red and yellow, respectively. The γ subunit of $TF_1$ in the binding dwell (PDB ID: 7L1Q) is shown in pink, and the γ subunit of $TF_1$ in the catalytic dwell (PDB ID: 7L1R) is shown in light blue. Both γ clashed with the α subunit facing $\beta_{80°}$ and with $\beta_{200°}$. Sites of steric clash are indicated by black stars.

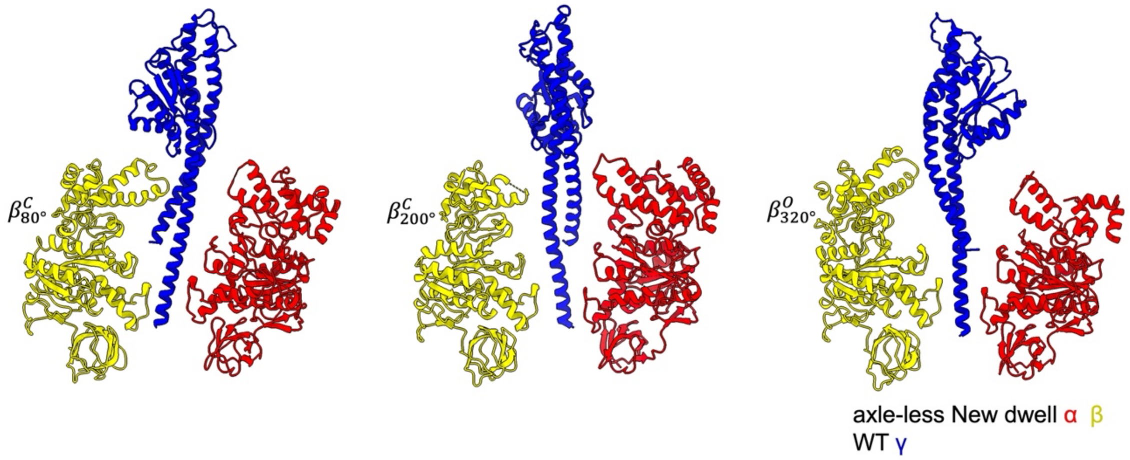


Fig. S14. Fitting of full-length γ in the new-dwell $\alpha_3\beta_3$ conformation of axle-less $TF_1$

WT $TF_1$ γ was fitted onto the $\alpha_3\beta_3$ ring of axle-less $TF_1$ in the new dwell. The γ structure represents an intermediate generated by linear interpolation between the binding dwell structure (PDB ID: 7L1Q) and the catalytic dwell structure (PDB ID: 7L1R). In the representation, the α, β, and γ subunits are shown in red, yellow, and blue, respectively.

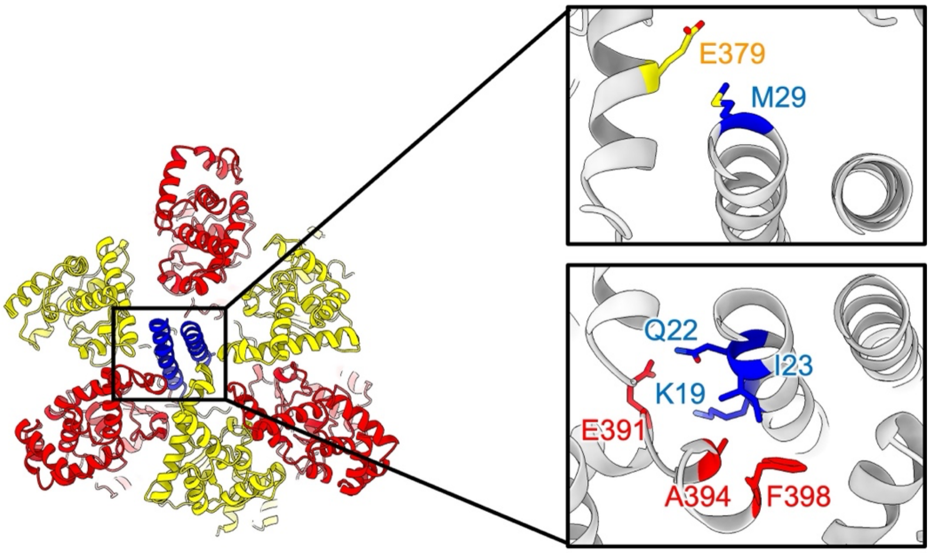


Fig. S15. New interactions between α/β and γ in the new dwell.

The α and β residues detected within 4.5 Å of γ in the new dwell, together with the corresponding γ residues, are shown. The upper-right panel shows the β–γ interactions, and the lower-right panel shows the α–γ interactions. The α, β, and γ subunits are colored red, yellow, and blue, respectively.

Table. S1. Cryo-EM data collection, refinement and validation statistics.

| | #1 Binding Dwell (EMD-69782) | #2 New Dwell (EMD-69783) | #3 Catalytic Dwell (EMD-69784) |
|---|---|---|---|
| **Data collection and processing** | | | |
| Magnification | 59,000x | 59,000x | 59,000x |
| Voltage (kV) | 300 | 300 | 300 |
| Electron exposure (e–/Å$^2$) | 74 | 74 | 74 |
| Defocus range ($\mu$m) | 0.5-1.5 | 0.5-1.5 | 0.5-1.5 |
| Pixel size (Å) | 0.84 | 0.84 | 0.84 |
| Symmetry imposed | C1 | C1 | C1 |
| Initial particle images (no.) | 608,830 | 852,517 | 845,464 |
| Final particle images (no.) | 422,075 | 464,363 | 415,389 |
| Map resolution (Å) masked FSC threshold 0.143 | 2.61 | 2.53 | 2.61 |
| Map resolution (Å) unmasked FSC threshold 0.143 | | | |
| Global resolution (Å) | 3.11 | 3.04 | 3.07 |
| Sphericity (threshold 0.5) | 0.790 | 0.803 | 0.790 |
| Map sharpening *B* factor (Å$^2$) | -61.1 | -60.6 | -62.0 |
